# Neural-symbolic hybrid model for myosin complex in cardiac ventriculum decodes structural bases for inheritable heart disease from its genetic encoding

**DOI:** 10.1101/2024.09.05.611508

**Authors:** Thomas P. Burghardt

## Abstract

**Background:** Human ventriculum myosin (βmys) powers contraction sometimes in complex with myosin binding protein C (MYBPC3). The latter regulates βmys activity and impacts overall cardiac function. Nonsynonymous single nucleotide variants (SNVs) change protein sequence in βmys or MYBPC3 causing inheritable heart diseases by affecting the βmys/MYBPC3 complex. Muscle genetics encode instructions for contraction informing native protein construction, functional integration, and inheritable disease impairment. A digital model decodes these instructions and evolves by continuously processing new information content from diverse data modalities in partnership with the human agent.

**Methods:** A general neural-network contraction model characterizes SNV impacts on human health. It rationalizes phenotype and pathogenicity assignment given the SNVs genetic characteristics and in this sense decodes βmys/MYBPC3 complex genetics and implicitly captures ventricular muscle functionality. When a SNV modified domain locates to an inter-protein contact in βmys/MYBPC3 it affects complex coordination. Domains involved, one in βmys and the other in MYBPC3, form coordinated domains (co-domains). Co-domains are bilateral implying potential for their SNV modification probabilities to respond jointly to a common perturbation to reveal their location. Human genetic diversity from the serial founder effect is the common systemic perturbation coupling co-domains that are mapped by a methodology called 2-dimensional correlation genetics (2D-CG).

**Results:** Interpreting the general neural-network contraction model output involves 2D-CG co-domain mapping that provides natural language expressed structural insights. It aligns machine-learned intelligence from the neural network model with human provided structural insight from the 2D-CG map, and other data from the literature, to form a neural-symbolic hybrid model integrating genetic and protein interaction data into a nascent digital twin. This process is the template for combining new information content from diverse data modalities into a digital model that can evolve. The nascent digital twin interprets SNV implications to discover disease mechanism, can evaluate potential remedies for efficacy, and does so without animal models.

**Highlights:** Neural-symbolic hybrid model decodes muscle genetics into contraction mechanisms And evolves in virtuous cycle *Optimize-Interpret-Revise-Repeat* aided by human partner Nascent digital twin unravels inheritable disease mechanism without animal models And estimates cardiac phenotype coupling strength to myosin thick-filament structures

## INTRODUCTION

Muscle proteins assembled in a sarcomere produce force and displacement by their coordinated action. In human ventriculum, cardiac myosin (βmys) forms the thick-filament and actin the thin-filament. The βmys ATPase motor projects outward from the thick-filament backbone to contact actin and repetitively convert ATP free energy into moving thin-filaments against resisting force. βmys and cardiac myosin binding protein C (MYBPC3) form a complex on the thick filament [2–4]. MYBPC3 regulates muscle activity through inter-protein contacts among domain sub-structures of the actin/βmys/MYBPC3 complex [5–7]. βmys /MYBPC3 inter-protein contacts are the focus here.

Nonsynonymous single nucleotide variants (SNVs) alter protein sequence in cardiac sarcomere proteins sometimes causing heart disease. A large collection of them contained in a SNV database informs basic SNV characteristics including four independent inputs that are known with certainty: sequence position, residue substitution, human population source, allele frequency, and two often unreported dependent outputs: phenotype and pathogenicity. A neural network models the causal links between these input SNV characteristics and heart disease phenotype and pathogenicity. It fills gaps in phenotype and pathogenicity assignment coverage [8, 9]. The 6-dimensional data point (6ddp) dataset (4 inputs and 2 outputs) describes functionality of a SNV modified human ventricular sarcomere. It is a system captured by a directed acyclic graph (DAG) relating molecular level inputs to outputs describing human health status (Model 6, **Figure 1**).

**Figure 1.**
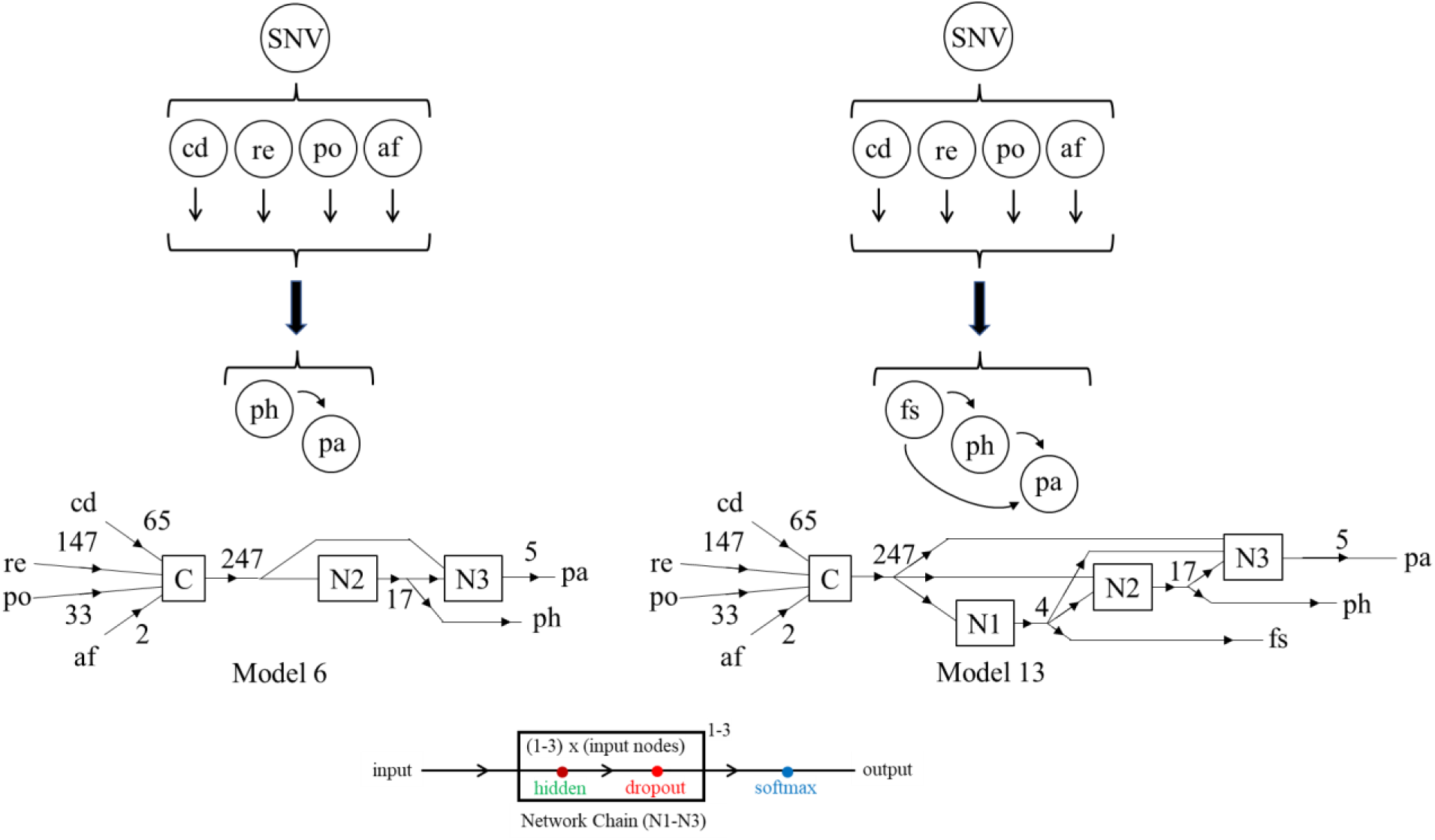
Directed acyclic graph (DAG) models for contraction (top row) and their neural network representation (middle and bottom rows). Contraction models are perturbed by SNVs located in the βmys/MYBPC3 complex as reflected in validating datasets (not shown) for Model 6 (left) or Model 13 (right). Model 6 was described earlier [1]. Model 13 introduces another output indicated by *fs* for thick *filament structure*. Inputs are functional protein domain (*cd*), residue substitution (*re*), human population source (*po*), and SNV frequency in the population (*af*). Outputs are filament structure (*fs*), disease phenotype (*ph*), and pathogenicity (*pa*). Arrows show direction of influence with no feedback. Neural networks (middle row) indicate inputs concatenated by the C-module (C). Numbers over arrows in the diagrams in the middle row indicate parameter number (e.g., 65 domains for the βmys/MYBPC3 complex at *cd*) that are input to Network Chains (N) depicted in more detail in the bottom row. The number of input nodes to each network chain is equal to the total inputs (247). Network chains have 1-3 hidden layers (indicated by the 1-3 superscript above the box in the net chain diagram) with total nodes numbering 1-3 times the number of input nodes. The dropout layer mitigates overfitting (along with another measure described in Methods) and the softmax layer converts the final output to digital form.

Model 6 is a general network contraction model interpreting molecular data in terms of a systemic outcome. It is a feed-forward neural network with subsets of fulfilled 6ddps, where input and outputs are known, forming validation datasets. Unfulfilled 6ddps, where one or both outputs are unknown, are fulfilled using the optimized neural network, creating the unfulfilled/fulfilled dataset. The combined dataset (fulfilled plus unfulfilled/fulfilled) has no unknows and, when interpreted probabilistically with a discrete Bayes network also based on the DAG, provides SNV probability for a functional domain location given phenotype and human population [8]. It supplies the information needed to identify phenotype specific, SNV impacted, co-domains in the βmys/MYBPC3 complex using a second method called 2-dimensional correlation genetics (2D-CG). 2D-CG locates SNV-affected inter-protein domain contacts within the βmys/MYBPC3 complex [10]. The 2D-CG map is the natural language translation of the machine intelligence contained in the Model 6 neural network general contraction model.

SNVs causing inheritable heart diseases including, dilated cardiomyopathy (DCM), familial hypertrophic cardiomyopathy (FHC), and left ventricle non-compaction (LVN), frequently target the βmys/MYBPC3 complex causing structural modification to functional domains that contain them [11]. βmys/MYBPC3 inter-protein contacts mediate contraction regulation in the native form and cause heart disease when modified by pathogenic SNV substitutions. 2D-CG is applicable to protein structural inquiries when genetic data specifies human population. Human populations evolve along distinct lines influencing their genetic divergence and selectively correlating protein residue sites coupled by their common involvement in an inheritable disease [1, 10, 12]. 2D-CG gives a natural-language and molecular level interpretation of what the genetics data implies about SNVs causing inheritable heart diseases.

2D-CG interpretation of Model 6 output gives structural substance to the implicit contraction mechanism residing in its trained neural network. Characteristic thick filament structural states emerge for DCM, FHC, and LVN phenotypes from the 2D-CG decoding when combined with thick filament cryo-EM structure data [2, 13] mainly related to DCM, and other data as follows:

i. DCM phenotype (composed of three separate phenotype categories indicated by 2 letter abbreviation *ds, d1,* and *dc*, see Supplementary Information (**SI) Figure S1**) is hypocontractile with a structure possibly akin to a hyper-relaxed sequestered [14], mavacamten-stabilized sequestered [2, 3], or mechanosensed-deactivated/off state of native thick filaments [4, 15, 16]. DCM inducing SNV modified myosins are more likely to be sequestered/off when compared to resting native myosins and less likely to actively develop force. The DCM βmys/MYBPC3 inter-protein contacts form the model for the sequestered/off state. They are surmised using *in vivo* human cardiac DCM 2D-CG maps [1, 10] and *in vitro* mavacamten-stabilized sequestered thick filament structure [2].
ii. FHC phenotype (*hx, h1, h4,* and *h8*, see **SI Figure S1**) reflects hypercontractile, destabilize sequestered myosin, disordered/on, or mechanosensed-activated (on) structural state because these SNV modified myosins are more likely disordered/on when compared to native and more likely to actively develop force. The FHC βmys/MYBPC3 inter-protein contacts form the model for the disordered/on state represented by the *in vivo* human cardiac FHC 2D-CG maps.
iii. LVN phenotype (*lc, lv,* and *lw*, see **SI Figure S1**) represents a third thick filament structural state where the SVN modified relaxed myosin heads have a specific but (for now) uncharacterized structural and functional state. The LVN βmys/MYBPC3 inter-protein contacts form the model for an uncharacterized, possibly new, structural state represented by the *in vivo* human cardiac LVN 2D-CG maps.
iv. Remaining phenotypes in the fulfilled dataset are several and represent a mixture of relaxed myosins in unknown thin-filament structural states. They are represented by the *in vivo* human cardiac NOT phenotypes 2D-CG maps. NOT phenotypes are members of the fulfilled plus unfulfilled/fulfilled 6ddps excluding DCM, FHC, and LVN phenotypes.

Filament states are defined by their equivalence with the broader phenotype categories (DCM, FHC, LVN, and NOT) in (i)-(iv). Natural language interpretations of their co-domain maps summarize findings. DCM co-domains link MYBPC3 C-terminal domains across 3 crown levels on two adjacent 430 Å myosin filament repeats of myosin dimers on the myosin thick filament surface. These SNV-enhanced interactions favor sequestered/off myosins shifting steady-state towards sequestered/off relaxed myosin heads. FHC co-domains link MYBPC3 N-terminal domains to 2-3 crown levels of myosin dimers. These SNV induced interactions favor disordered/on myosins shifting steady-state towards disordered/on myosin heads. LVN co-domains link MYBPC3 N-through C-terminal domains to multiple crown levels of relaxed myosin dimers. It implies a comprehensive alteration of native βmys/MYBPC3 inter-protein contacts. NOT co-domains are a heterogeneous mixture of relaxed myosin structural states that reveal their in-common interactions between MYBPC3 and myosin thick-filament backbone.

What genetics data implies about SNVs causing inheritable heart diseases is summarized by these natural-language molecular interpretations. Co-domain maps for each phenotype (DCM, FHC, and LVN) identifies unique bi-molecular interaction footprints reflecting divergent disease mechanisms. They identify critical and possibly dominant myosin thick filament structures among the DCM, FHC, and LVN phenotypes leading to a new general network contraction model following from Model 6 but explicitly incorporating a dynamic thick filament structure parameter (*fs*) in a follow up model (Model 13, see **Figure 1**). It merges machine-learning and human insight into a neural-symbolic hybrid model for the structural basis of disease, enlarging 6ddps to 7ddps with *fs* joining dependent outputs phenotype (*ph*) and pathogenicity (*pa*). The hybrid model database (both genetic and human-insight based) develops with time and will influence successor contraction models incorporating greater complexity involving more interacting proteins into the machinery.

The neural network machine-learning with human-insight merger has potential to accelerate application of more comprehensive models for contraction to understand and manage inheritable heart disease using a human heart digital twin. The nascent digital twin introduced here interprets SNV implications to discover disease mechanism, can evaluate potential remedies for efficacy, and does so without animal models [17].

## RESULTS

### βmys/MYBPC3 co-domains

**SI Figure S2** indicates homology (βmys) and linearized models (βmys and MYBPC3), many of their structural/functional domain locations, and two letter abbreviations for most domains involved in co-domains. **SI Figure S3** lists all domains and their sequence assignment in the βmys/MYBPC3 complex.

### General network contraction Model 6

**Figure 2 panels a-c** show Model 6 2D-CG maps identifying co-domains linking βmys with MYBPC3 in the βmys/MYBPC3 complex for DCM, FHC, and LVN phenotypes. Maps are constructed as described in METHODS section **2D correlation intensity and significance** with eq. 11 and as described [1]. Each pixel corresponds to a single co-domain pair and grayscale representation proportional to their SNV probability amplitude product. **Figure 2 panels a and b** for DCM and FHC phenotypes are identical to earlier work (**Figure 4** in [1]) while **panel c** for LVN phenotypes is revised by co-domain re-assignment from {*lx,m5*} and {*lx,en*} to {*l6,rn*} and {*lx,rn*}, respectively. The revision reflects fluctuation in neural network optimization that involves generating scenarios differing by hidden layer width, depth of layers, node weights and biases optimization, and by training data randomly selected from the validation data set (see METHODS: **Neural network training and validation**). It introduces rank score fluctuation that shifted the {lx,m5} and {lx,en} co-domains from positions 5 and 6 to 11 and 13 in the current simulations. Prior 6 most significant co-domains are 14 standard deviations from mean compared to 17 for the current 6 implying improved optimization in this round. Rank score fluctuations affected only the LVN phenotype probably because of its overall lower standard deviation contrast among SNV detected co-domains.

**Figure 2.**
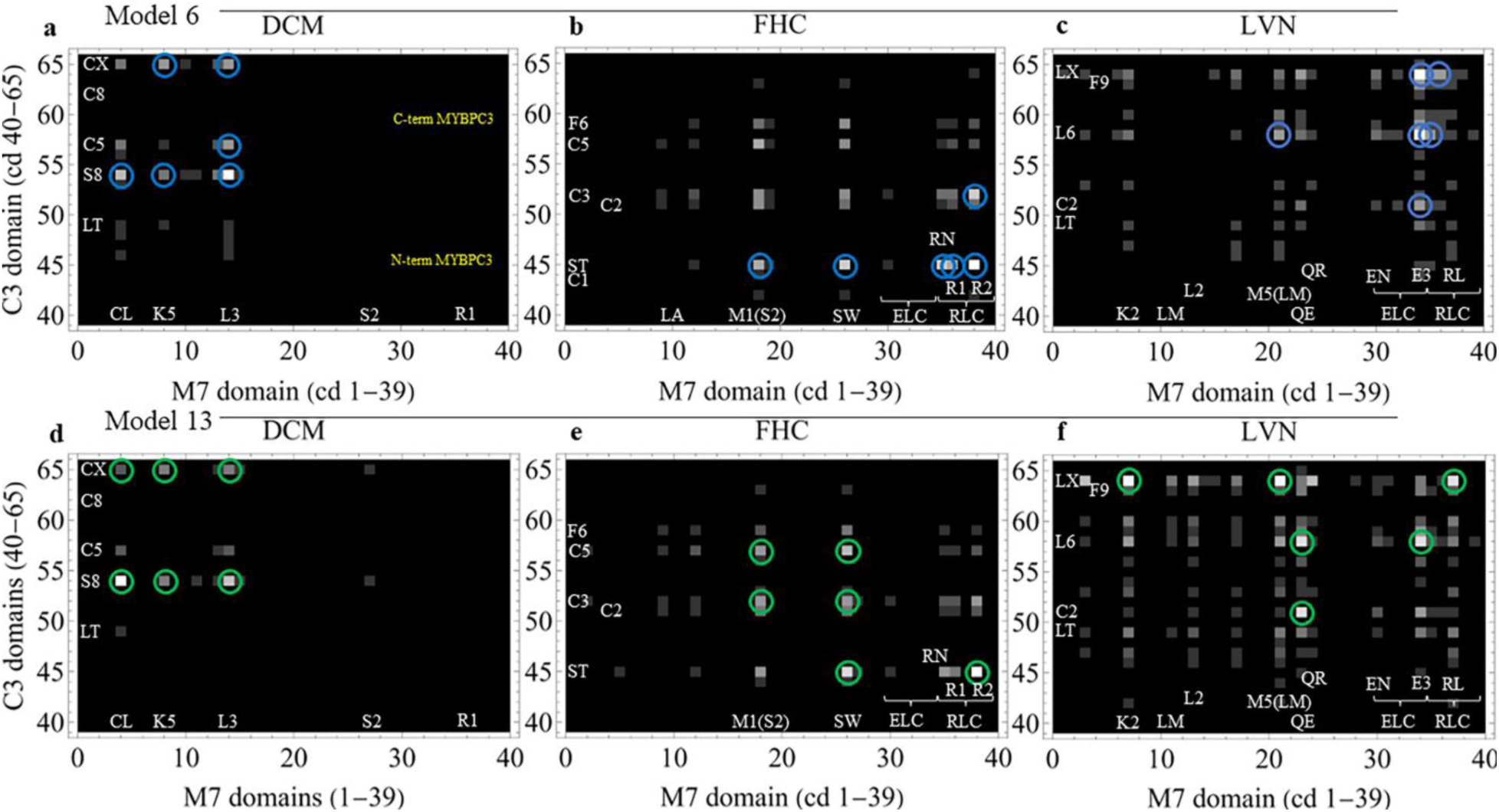
Model 6 (**panels a-c**) or Model 13 (**panels d-f**) 2D-CG cross-correlated SNV probability maps linking co-domains in the βmys/MYBPC3 complex for DCM, FHC and LVN phenotypes. (**panel a**) Cross-correlate intensity indicated by the grey scale (black=0) for DCM. Axes have βmys (M7) domains (1-39 along abscissa) and MYBPC3 (C3) domains (40-65 along ordinate). Selected domains forming co-domains are labeled with their two letter codes. Domain visualization, indexing, and two letter codes are from **SI Figures S2-S3**. Blue circles indicate the 6 most significant co-domains. (**panels b-c**) Notation identical to **panel a** but for FHC and LVN phenotypes. (**panels d-f**) Notation identical to **panels a-c** except with green circles indicating the 6 most significant co-domains for Model 13.

Co-domain probability amplitudes are distributed normally about zero. Outliers include the 6 most outstanding indicated by blue circles in the maps. They fall >107, >118, and >17 standard deviations from the mean for the DCM, FHC, and LVN phenotypes, respectively. Co-domain outliers identify the most certain coordinating interactions between βmys and MYBPC3. Co-domains link βmys motor domain and lever-arm bound light chains with MYBPC3 N-terminal segment (c0-c5) or C-terminal segment (c5-cx) domains, respectively. Inspection of **Figure 2** 2D-CG maps show how the co-domain distributions differentiate these three phenotypes.

**Figure 3a** has a cartoon model of human ventriculum cardiac myosin thick filament based on the cryo-EM structure from Dutta et al. [2]. Thick filament myosin has sequestered dimer heads forming the interacting-heads motif (IHM) containing free (blue) and blocked (red) motor domains, lever-arm bound light chains ELC (green) and RLC (orange), and tail domains (*s2* and *lm*, light grey) collectively forming the thick filament backbone. MYBPC3 has domains *c0*-*cx* connected by linker domains (dark grey) lying on the thick filament surface and contacting multiple sequestered myosin dimer heads. The βmys IHM dimers lie on the thick filament surface in (crown) levels D (CrD), T (CrT), and H (CrH) forming a 430 Å repeating structural feature. In the cartoon, M-line is nearer the MyBPC3 C-terminus at *cx*. βmys dimers in IHM’s link to each other at contacts between the CrD blocked head ELC and CrT free head motor domain, and, between the CrT blocked head RLC and CrH free head motor domain. The leftmost CrD represents an adjacent 430 Å repeat. It is displaced from CrH by a larger azimuthal rotation (32 degrees) compared to the azimuthal separation between CrT and CrH (16 degrees).

**Figure 3.**
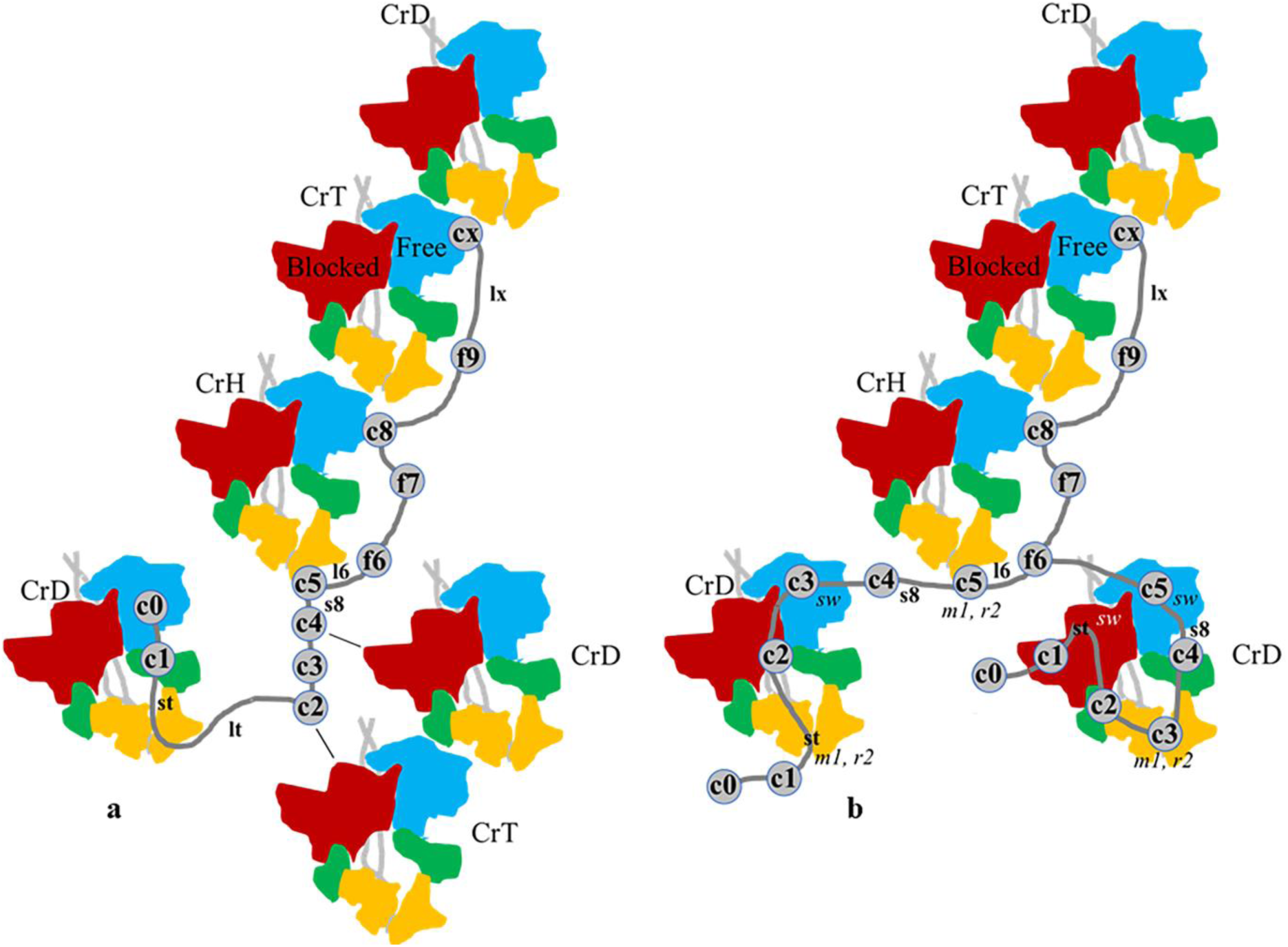
(**a**) Cartoons representing sequestered/off βmys dimers in crown levels CrD, CrT, and CrH forming the 430 Å-long repeating quasihelical strand on the thick filament surface from interpreting a cryo-EM reconstruction [2]. Myosin heads are in blocked (red) or free (blue) conformations with light chains ELC (green), RLC (orange), and myosin S2 (light grey). MYBPC3 domains are numbered as in **SI Figure S2** and linkers shown in dark gray. The adjacent CrD-CrT partial strand on the lower right shows weaker βmys/MYBPC3 co-domains (black lines) involving MYBPC3 domains *c2* and *c4* and blocked head motor domains in CrT and CrD. Partial strand containing CrD on the left indicates a proposed transient interaction with the MYBPC3 N-terminus that competes with a bound actin option (not shown). (**b**) Proposed disordered/on co-domains from interpreting the Model 13 FHC 2D-CG map shown in Figure 2e and as described in Results.

**Figure 3a** indicates βmys/MYBPC3 contacts by overlapping protein domains at CrT(Free head motor domain)/*cx*, CrH(Free head motor domain)/*c8*, and CrH(RLC)/c5. The CrT(Free head motor)/*cx* and CrH(Free head motor)/*c8* contacts form the signature super-relaxed (SRX) βmys/MYBPC3 complex by implying ≥2/3 of free heads are sequestered over the 430 Å repeats involving MYBPC3. MYBPC3 domains *c2* through *cx* also maintain contacts with the myosin filament backbone. MYBPC3 domains *c0*, *c1*, and *c1-c2* linker (*lt*) apparently form transient association with the lowest level CrD(Free head) as shown at the lower left of **Figure 3a**, or actin [18] in the native environment. MYBPC3 N-terminal domains *c0-c4* form weak (*c2-c4*) and weaker (*c0-c1*) thick filament links. Lower contrast cryo-EM data suggests *c2* binds CrT(Blocked head motor) and *c4* binds CrD(Blocked head motor) adjacent myosin head domains shown on the lower right of **Figure 3a**. The **Figure 3a** cartoon is a representation for the relaxed sequestered/off filament structural state. Co-domain formation sites identified with DCM 2D-CG findings refer to this diagram but equivalence between the actual relaxed sequestered/off filament structural state and SRX is inconclusive [2–4, 16].

### Neural-symbolic contraction model

A general network contraction model like Model 6, together with human interpretation of its 2D-CG map and findings from the literature, merges machine learning with human insight providing incentive and means to introduce and assign a thick filament structure state parameter, *fs*, in a follow-up model that is Model 13 (**Figure 1**). Model 13 is a neural-symbolic hybrid describing the structural basis of disease for DCM, FHC, and LVN phenotypes and with dependent filament state *fs* joining the other dependent parameters, phenotype (*ph*) and pathogenicity (*pa*).

Filament state *fs* assumes one of 5 values {*sr, dr, xr, gr, uk*}, defined descriptively by sequestered/off-relaxed (*sr*), disordered/on-relaxed (*dr*), other-relaxed (*xr*), generic-relaxed (*gr*), and unknown (*uk*). Its assignment implies location and impact of residues in co-domains mediating MYBPC3 regulation of βmys. It expands the 6ddp data set to the 7ddp format {*cd, re, po, af; ph, pa*}→{*cd, re, po, af; fs, ph, pa*} where the semi-colon separates independent (left) from dependent (right) variables, and with unknown (*uk*) initially filling in all *fs* positions. Co-domains for DCM, FHC, and LVN phenotypes indicated by Model 6 are plotted in **Figure 2** with six most significant identified by blue circles. These data correspond to ordered pairs of domains, one from βmys and the second from MYBPC3, identifying inter-protein contacts impacting thick filament structure. These data and other information from the literature constrain Model 13 optimization by influencing SNV assignment of *fs* in fulfilled 7ddps. The process is iterative, adaptive, and illustrated in **SI Figure S4**.

It is presumed that βmys/MYBPC3 co-domain contacts influence thick filament structure related to myosin contractility regulation. 2D-CG assignment of βmys/MYBPC3 co-domains for each inheritable disease (**Figure 2**) is the primary source for filament state assignment. Thick filament stability conferring contacts surmised from cryo-EM data pertains exclusively to the mavacamten-stabilized SRX state and its structural insights are used accordingly. Other information about co-domain interactions is from the literature as described below.

Case 1. Super-Sequestered/off (*fs* = *sr*)

### βmys/MYBPC3 co-domains stabilizing sequestered/off myosin heads implicated by cryo-EM

Human cardiac ventriculum myosin thick filament structure in mavacamten-stabilized SRX state has free head motor domains in myosin dimers CrT and CrH contacting MYBPC3 C-terminal domains *cx* and *c8*, respectively. The CrH free head RLC also contacts *c5*. Lower contrast, transient βmys/MYBPC3 co-domain contacts involve blocked head motor domains from adjacent CrT and CrD myosins with MYBPC3 *c2* and *c4*, and, association of free head CrD with *c0* and *c1* (**Figure 3a**) [2]. Thick filament structure (PDB: 8G4L) implies βmys domains, adapted to the 2D-CG domain mapping preferences in **SI Figure S3**, have βmys free head motor domains *cL* and *bh* forming co-domains with both *cx* and *c8* while CrH free head RLC domains *r1* and *rL* form co-domains with *c5*. Domain *cL* is the C-Loop [19, 20] and *bh* the blocked head converter binding site [21]. These domains from the βmys/MYBPC3 complex form the set {*cL, bh, r1, rL*, /, *cx, c8, c5*}.

### βmys/MYBPC3 co-domains stabilizing sequestered/off myosin heads implicated by 2D-CG

Apparent differences in βmys/MYBPC3 co-domain contacts over the DCM, FHC, and LVN phenotypes (compare **Figures 2a-2c**) indicate the SNV induced alterations in βmys/MYBPC3 interactions. It manifests in DCM or FHC cases with super-sequestered/off (Case 1) or super-disordered/on (Case 2) shifts from native off SRX vs on DRX populations [22–24], or, mechanosensed-off vs -on populations [16]. The “super-“ prefix for Case 1 indicates the presumption that DCM or FHC SNVs shift the native complex myosin ensemble into more uniform sequestered/off or disordered/on ensemble states.

Model 6 2D-CG maps from DCM causing SNVs (**Figure 2a** and blue circles) implicate CrT(Free head motor at *k5* and *L3*)/*cx*, CrH(Free head motor at *L3*)/*c5*, and CrD(Blocked head motor at *cL*, *k5*, and *L3*)/*c4* (*s8* is a phospho-threonine within *c4*) co-domains in general agreement with expectations from the cryo-EM except for the MYBPC3 interaction with CrH (cryo-EM identifies CrH(Free head)/*c8* and CrH(Free head RLC)/*c5* co-domains). The 2D-CG identified six most significant βmys/MYBPC3 co-domains are indicated in **Figure 2a** by blue circles. βmys domains include C-loop (*cL*), 50 kDa motor domain proteolytic fragment (*k5*), and loop 3 (*L3*). All are or contain actin binding sites while *k5* also includes portions of the active site and none of them involve RLC. That βmys RLC and MYBPC3 *c8* fail to appear within the 6 most significant DCM co-domain list from Model 6 2D-CG data is discussed later in RESULTS when considering the SNV distribution over sites in the βmys/MYBPC3 complex. Domains from complexed βmys/MYBPC3 forming the super-sequestered/off myosin heads from DCM 2D-CG data form the set {*cL, k5, L3*, /, *cx, c5, c4(s8)*}. Overlaps with cryo-EM identified domains include {*cL, /,cx, c5*}. Combining 2D-CG and cryo-EM identified co-domains gives the super-set {*cL, bh, k5, L3, r1, rL,* /, *cx, c8, c5, c4(s8)*}.

### βmys and MYBPC3 domains instigating super-sequestered/off myosin heads form the new fulfilled 7ddps

SNVs participating in βmys/MYBPC3 co-domains identified in Case 1 from the fulfilled 6ddp dataset and not assigned with FHC or LVN phenotypes are assigned the filament structure *sr* and become members of the new fulfilled 7ddp dataset.

SNVs participating in the βmys/MYBPC3 co-domains identified in Case 1 from the unfulfilled 6ddp dataset and not assigned with FHC or LVN phenotypes are likewise assigned the filament structure *sr* and form the unfulfilled 7ddp dataset. These assignments in the unfulfilled 6ddps (converting them to unfulfilled 7ddps) do not impact subsequent Model 13 network optimization (that depends only on fulfilled 7ddps), but they will influence other quantities discussed subsequently that depend on the fulfilled plus unfulfilled/fulfilled dataset, i.e., all members of the 7ddp dataset.

### Summarizing assignments

The βmys/MYBPC3 domain set associated with cryo-EM structure (PDB 8GL4), DCM SNVs, or both, contains elements {*cL, bh, k5, L3, r1, rL,* /, *cx, c8, c5, c4(s8)*} and obtain filament state assignment *sr* in the 7ddps as described in Case 1. It implies interactions between the MYBPC3 C-terminus (*c4*-*cx*) and three myosin crown levels, CrT(Free head motor), CrH(Free head motor), and CrD(Blocked head motor). Interactions considered but left unassigned, including the CrT(Blocked head motor)/*c2* and CrD(Free head)/(*c0-c1*), likewise associate with filament states other than *sr* or lack supporting structural data.

Case 2. Super-Disordered/on (*fs* = *dr*)

### βmys/MYBPC3 co-domains stabilizing disordered/on myosin heads implicated by 2D-CG

**Figure 2b** shows Model 6 FHC 2D-CG cross-correlates between: (*i*) *st* (MYBPC3 N-terminal serine 2 in the regulatory domain between *c1* and *c2*) and RLC (*rn*, *r1*, and *r2*), (*ii*) *st* and switch 2 helix (*sw*) in the myosin motor domain, (*iii*) *st* and subfragment 2 (*s2*) at *m1*, and (*iv*) *c3* and RLC (*r2*). These 6 most significant 2D-CG cross-correlates associated with the FHC phenotype are consistent with relaxed disordered/on state especially when MYBPC3 is phosphorylated at *st* [5]. Lower significance co-domains within the 11 most significant imply cross-correlation of MYBPC3 domains *c2*, *c5* and *f6* are visibly notable, see **Figure 2b**. Mentioned βmys/MYBPC3 interactions implicate MYBPC3 N-terminal domains {*st, c2, c3, c5, f6*} and βmys domains {*m1, rn, r1, r2, sw*}.

Case 1 and Case 2 domains overlap at *r1* and *c5*. When there is overlap, Cases 1-4 identified domains are prioritized by significance in the order Case 1 > Case 2 > Case 3 > Case 4 unless phenotype is assigned FHC or LVN, then phenotype decides SNV filament structure assignment. Case 2 identified βmys/MYBPC3 domains are the set {*m1, rn*, *r1, r2, sw, /, st, c2, c3, c5, f6*} from the 11 most significant co-domains for FHC phenotypes indicated by **Figure 2b** in 6 blue circles and also including 3 visibly notable co-domains at *m1*/*c2*, *m1*/*c5*, and *sw*/*f6*. **Figure 3b** show proposed *super-disordered/on* state structures consistent with Case 2 co-domains.

### βmys and MYBPC3 domains instigating super-disordered/on myosin heads incorporated into the new fulfilled 7ddps

SNVs participating in the βmys/MYBPC3 co-domains identified in Case 2 from the fulfilled 6ddp dataset, and not assigned with LVN phenotype, are assigned filament structure *dr* and become members of the new fulfilled 7ddp dataset.

SNVs participating in the βmys/MYBPC3 co-domains identified in Case 2 from the unfulfilled 6ddp dataset and not assigned with LVN class phenotypes (*lc*, *lv*, *lw*), are likewise assigned filament structure *dr* and form the unfulfilled 7ddp dataset.

### Summarizing assignments

Case 2 βmys/MYBPC3 domain set {*m1, rn*, *r1, r2, sw, /, st, c2, c3, c5, f6*} identifies the MYBPC3 N-terminus (*st*-*f6*). The βmys crown levels CrH at RLC domains and CrD at free head myosin (**Figure 3b left side**), or, free and blocked head myosins (**Figure 3b right side**).

Case 3. Other-relaxed filament state (*fs* = *xr*)

### βmys and MYBPC3 domains identified with other-relaxed thick filament structure are incorporated into fulfilled 7ddps

*β*mys and MYBPC3 domains identified by LVN SNVs forming the 12 most significant co-domains include {*e3, k2, m5, qe, /, c2, f9, L6, Lx*}. Case 2 and Case 3 domains overlap at *c2* is managed as already described. Furthermore, remaining 6ddps with *fs* = *uk*, after consideration in Cases 1-3, are assigned exclusively by phenotype with DCM phenotypes getting *fs* = *sr*, FHC phenotypes *dr*, and LVN phenotypes *xr*.

SNVs participating in the βmys/MYBPC3 co-domains identified in Case 3 from the unfulfilled 6ddp dataset are assigned the other relaxed thick filament structure (*xr* except as noted above for *c2*).

Case 4. Generic-relaxed filament state (*fs* = *gr*)

Remaining unassigned filament structure parameters in fulfilled 6ddps are upconverted to fulfilled 7ddps by their assignment to the structurally unspecified generic-relaxed state (*gr*). It maintains column length of the fulfilled 7ddp dataset matrix (4092 x 7) equal to its 6ddp counterpart (4092 x 6).

A final unfulfilled 7ddp dataset (where at least one dependent parameter in every row is unknown) does not impact network training for the optimized neural network. However, the optimized neural network computes all unfulfilled dependent parameters in the unfulfilled 7ddp dataset providing the unfulfilled/fulfilled dataset. The unfulfilled/fulfilled dataset supplies information needed for computing conditional probability tables (CPTs, METHODS eq. 1) used to construct 2D-CG maps and thick filament structure basis (METHODS eqs. 12-13).

## Model 13

Model 13 neural network optimization and 2D-CG map construction from genetics data follows the protocol used previously with Model 6 [10] and as described in METHODS (sections **Neural network training and validation** and **2D correlation intensity and significance**). **Figure 2 panels d-f** indicate maps for the Model 13 2D-CG co-domains linking βmys with MYBPC3 for DCM, FHC, and LVN phenotypes. Each pixel in a map corresponds to one co-domain pair. Co-domain probabilities are distributed normally about zero with amplitudes falling >122, >111, and >19 standard deviations from their mean value for DCM, FHC, and LVN phenotypes, respectively. Their significance equivalent to that for Model 6. Outlier co-domains identify those most likely implicated in βmys/MYBPC3 interactions. The LVN co-domains are less significant than their DCM and FHC counterparts. Maps in **Figure 2** compare Model 6 and Model 13 results. They are qualitatively similar with intensity significance variation indicated by re-arrangement of some most significant co-domains (compare blue and green circles for equal phenotypes).

### Model 13 DCM and FHC 2D-CG maps

Model 6 and Model 13 2D-CG maps are similar for DCM (compare **Figures 2a and 2d**) and FHC (**Figures 2b and 2e**) implying the neural network and symbolic aspects of the hybrid model map do not conflict. Model 13 DCM 2D-CG has modestly higher emphasis on co-domains linking βmys with MYBPC3 C-terminus at *cx* while FHC 2D-CG has lower relative emphasis on co-domains linking βmys with MYBPC3 N-terminus at *st*.

Cryo-EM data indicates mavacamten-sequestered myosin free heads form co-domains CrT/*cx* and CrH/*c8* where the βmys actin and blocked-head-converter-binding domains (*cL* and *bh*) interact with both MYBPC3 domains *cx* and *c8*. Furthermore, RLC (*r1* and *rL*) from CrH forms co-domains with *c5*, and blocked heads in CrD/*c4* and CrT/*c2* (myosin sites unspecified) form weaker co-domains (see **Figure 3a**). This approximately parallels the Model 13 2D-CG footprints for DCM phenotypes (**Figures 2d**) linking sequestered myosins over three βmys dimer crown levels from free head CrT/*cx* to blocked head CrD/*c4(s8)* (**Figure 3a**). DCM involved myosin heads have actin binding domains (*cL*, *k5*, and *L3*) interacting with MYBPC3 *cx* and *c4* for the super-sequestered/off (*sr*) filament structure state. DCM 2D-CD data does not indicate the CrH/*c5* interaction involving RLC suggesting either that a SNV located there promotes myosin disorder and causes FHC or that the super-sequestered/off state is not equivalent to SRX [16]. The former is plausible since just 12 DCM and no LVN inducing SNVs target RLC while 499 FHC inducing SNVs locate to RLC in the SNV database (**Figure 4**).

**Figure 4.**
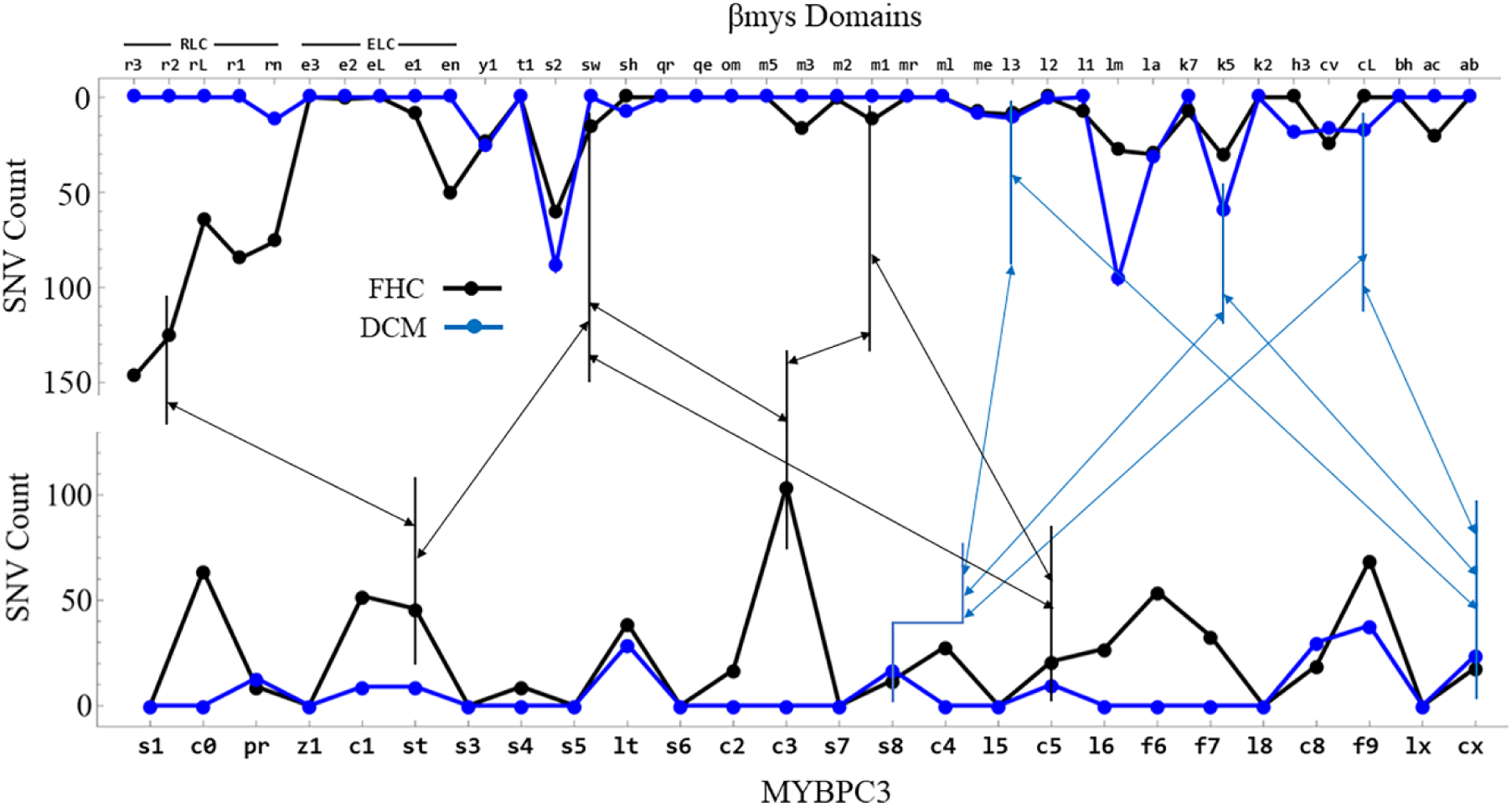
SNV count from the 7ddp dataset entries for known phenotypes, distributed over βmys and MYBPC3 domains for FHC (large black data points connected by thick black lines) and DCM (large blue data points connected by thick blue lines) defined by the protein domain sequence in **SI Figure S3**. Thin bidirectional arrows connect most-significant co-domains identified with Model 13 for FHC (black) and DCM (blue). The site identified as *s8* (phosphothreonine 8) in MYBPC3 is physically inside *c4* and closer to *c5* than *c3* while its location in the domain listing is between c3 and c4. The domain order in this figure follows the domain listing in **SI Figure S3** while the elbow connector better indicates the physical location of *s8*. Co-domains segregate into MYBPC3 N-terminal domains for FHC and C-terminal domains for DCM. FHC βmys co-domains favor RLC and the vicinity on the lever-arm (*r2, m1*) or motor domain (*sw*) while DCM βmys co-domains favor actin binding domains (*cL, k5, L3*) or active site (*k5*).

Comparing myosin binding sites for MYBPC3 in the cryo-EM (*cL* and *bh*) vs DCM 2D-CG (*cL*, *k5*, and *L3*) data indicates they overlap except at *L3* (an actin binding site on myosin). Mavacamten-sequestered myosin heads exclude the *L3*/MYBPC3 interaction suggesting *L3* forms a co-domain with MYBPC3 promoting a hyper-sequestered/off rather than sequestered/off filament state, or, the inequivalence of SRX and sequestered/off.

FHC phenotypes have relaxed myosin head domains detached from the thick filament surface in the super-disordered/on relaxed state. MYBPC3 continues to interact with myosin heads while the extreme C-terminus anchors it to *lm* [5, 25]. 2D-CG does not detect the latter co-domains probably because they lack SNVs (discussed below involving **Figure 4**). FHC 2D-CG detects βmys/MYBPC3 co-domains with the MYBPC3 N-terminal domains *c5-c0* that are mapped in **Figure 2e**. The map identifies interaction sites consistent with the two configurations shown in **Figure 3b**. Myosin head disorder is not indicated in the figure (**3b**) for simplicity. The 2D-CG map shows βmys at {*m1, sw*, and *r2*} potentially interacting simultaneously with MYBPC3 at {*c5, c3*, and *st*}. βmys *m1* and *r2* domains are spatially close making them amenable to simultaneous co-domain formation with MYBPC3, however, βmys *sw* in the motor domain is unlikely to join them while in the disordered head configuration (more likely in sequestered myosin at the CrT-CrH junction where RLC, s2, and free motor domain merge locations). The FHC 2D-CG rather implies that these domains form independent point contacts such as those identified in **Figure 3b** in the two scenarios involving adjacent 430 Å myosin filament repeats. There a flexible and mobile MYBPC3 N-terminus has *c3* (right side) or *st* (left side) contacting βmys *m1* and *r2*, while c5 (right) or c3 (left) contact βmys *sw*. The FHC 2D-CG map is consistent with other structural scenarios not explored here and all are probably transient.

Cryo-electron tomography on relaxed human cardiac myofibrils indicated the 430 Å myosin crowns repeats (CrD-CrT-CrH) provide the different myosin activation levels (relaxed-sequestered/off, relaxed-disordered/on, active) [4]. The scenario depicted in **Figure 3b**, derived from *in vivo* human data, has the MYBPC3 N-terminus shared with a disordered/on level (right) or an active level when *c0* and *c1* bind the thin-filament (left).

**Figure 4** shows SNV count distributions for DCM and FHC phenotypes for MYBPC3 (bottom) and βmys (top) domains and with Model 13 co-domains linking βmys with MYBPC3 identified by the two headed arrows. DCM SNV count distribution (thick blue lines) has 12 SNVs in RLC (top thick blue line, all in *rn*) while FHC SNV count distribution has 499 SNVs in RLC (top thick black line). Apparently, RLC SNVs usually cause FHC or another phenotype but not DCM possibly explaining the lack of CrH(Free head RLC)/*c5* co-domains in DCM 2D-CG (**Figure 2a & 2d**). Cardiac ventricular myosin RLC phosphorylation regulates myosin sequestered state occupation [14, 26, 27]. It suggests a pivotal significance for RLC in stabilizing sequestered/off myosin. Furthermore, FHC SNV count distribution has no SNVs in *cL* and *bh* (top thick black line) explaining the absence of co-domains combining them with *cx* or *c8* in FHC 2D-CG. SNVs in the *cx* or *c8* positions could destabilize MYBPC3 thick filament binding in the disordered/on state initiating a different phenotype.

Mavacamten sequestered βmys has MYBPC3 *c8* contacting CrH(Free head motor) in the blocked head converter binding site (*bh*, i.e., site on free head where blocked head converter domain *(cv)* binds), and C-loop (*cL*) actin binding site [2]. The Model 6 and Model 13 DCM 2D-CG maps register the *cL/c8* co-domain with low significance. As such, it is not visible in **Figures 2a** or **2d** relative to higher significance co-domains although both sites in the *cL/c8* co-domain host SNVs (top/bottom thick blue lines, **Figure 4**). The lower significance indicates lower correlation of these co-domain partners over the genetic divergence perturbation. Additional discussion of this exception for *c8* appears in a subsequent section.

### Model 13 LVN 2D-CG map

Model 6 and Model 13 LVN 2D-CG maps diverge more radically that those from DCM and FHC phenotypes. Rearrangement of the most-significant co-domains affects 5 of 6 assignments. The lower significance of all the LVN co-domains suggests normal modeling variations cause most significant co-domain site shifting. Model 13 2D-CG findings for LVN phenotype (**Figure 2f**) contrast with the DCM and FHC phenotypes. LVN has co-domains involving MYBPC3 C-terminus at *lx* and βmys *k2* (free head active site and lever arm), *m5* (in *lm*) binding site for *cx* [28]), and *rL* (RLC) implying a loosely bound MYBPC3 C-terminus at CrT (see **Figure 3a**). Another co-domain involves the central part of MYBPC3 at *l6* with βmys ELC (*e3*) and the ELC IQ domain on the lever arm (*qe*) implying a loosely bound MYBPC3 now at CrH (see **Figure 3a**). Finally, the MYBPC3 N-terminus at *c2* with βmys *qe* in CrD from an adjacent 430 Å myosin filament repeat like that in **Figure 3b** (right).

**Figure 5** shows SNV count distributions for LVN and FHC phenotypes over MYBPC3 (bottom) and βmys (top) domains with Model 13 co-domains identified by the two headed arrows. FHC SNV count distribution is identical to that in **Figure 4**. Here MYBPC3 contacts with βmys appear spontaneously without a physical LVN associated βmys SNV in the 7ddp data set (top thick blue line in **Figure 5**). The SNV modified sites are real but their association with LVN was made explicit only upon completion of unfulfilled 7ddps using the Model 13 neural-symbolic hybrid construct. Similarly, Model 13 anticipated specific RLC stabilization of the sequestered/off state by apparently ruling out a DCM associated RLC SNV when completing the unfulfilled 7ddps (**Figure 2d**).

**Figure 5.**
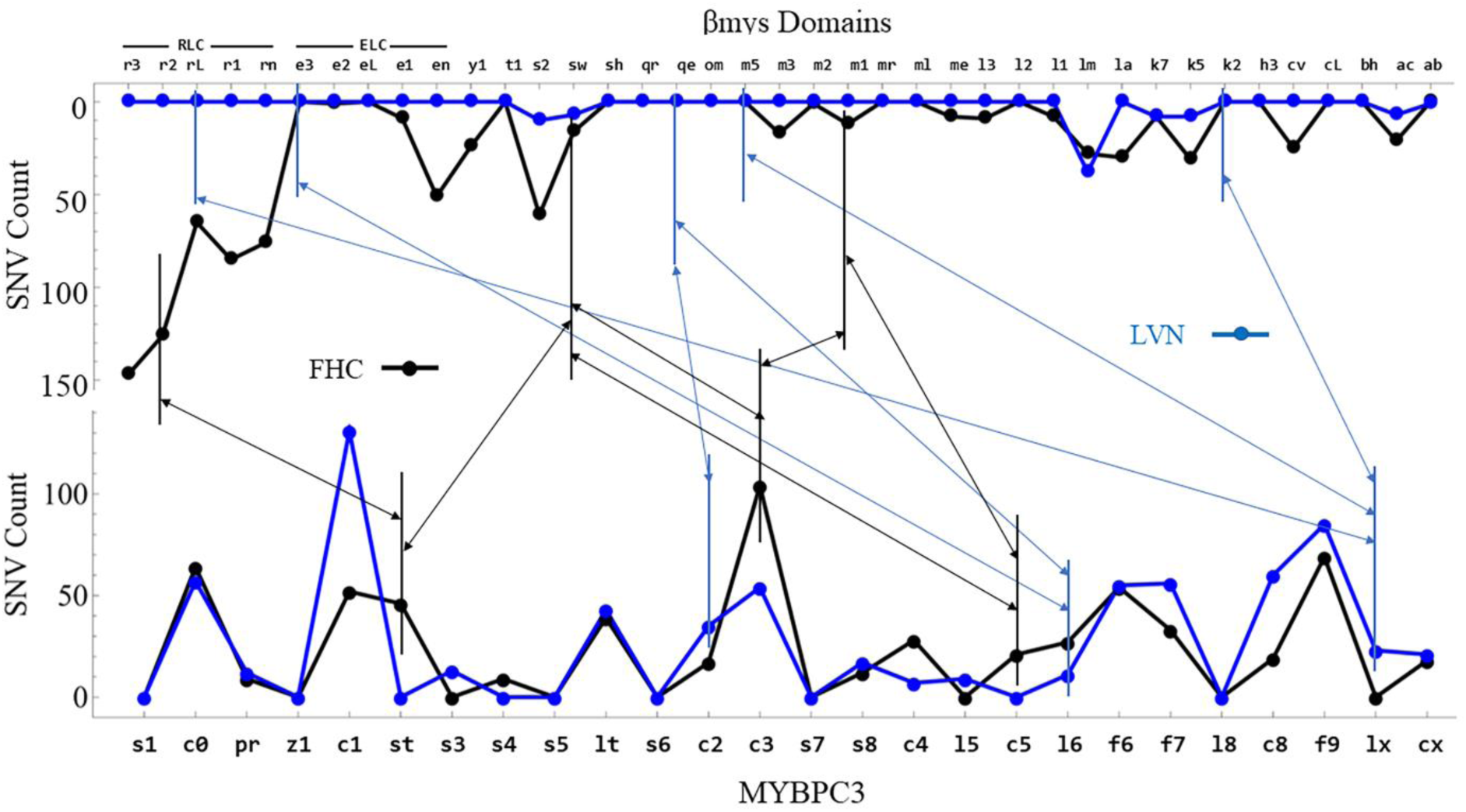
SNV count from the 7ddp dataset entries for known phenotype, distributed over βmys and MYBPC3 domains for FHC (large black data points connected by thick black lines) and LVN (large blue data points connected by thick blue lines) defined by the protein domain sequence in **SI Figure S3**. Thin bidirectional arrows connect most-significant co-domains identified by Model 13 in FHC (black) and LVN (blue). Most LVN co-domains in βmys have zero count in the dataset where just 7 domains, *ab*, *ac, k5, k7, lm, sw,* and *s2* host SNVs. Model 13 predicted βmys SNVs for the LVN phenotype form the set of most-significant co-domains. The FHC SNV count distribution in identical to that in Figure 4.

### NOT phenotypes

βmys or MYBPC3 domains assigned generic filament state structure (*gr*) form a fulfilled 7ddp subset for a diverse collection of phenotypes not including DCM, FHC, and LVN called the NOT phenotypes. NOT phenotypes have the 2D-CG map in **Figure 6a** indicating most-significant co-domains between MYBPC3 sites along its full length, from the N-terminus at a phosphorylatable serine (*s1*) to *c8*, with myosin filament backbone sites in *s2* and *lm*. Their most significant co-domains fall into the extreme N-terminus end of MYBPC3 (in the *c0* and *c1* domains) unlike the other phenotype groups. Most significant NOT co-domains do not preferentially engage the myosin motor domain suggesting they favor relaxed disordered/on filament state.

**Figure 6.**
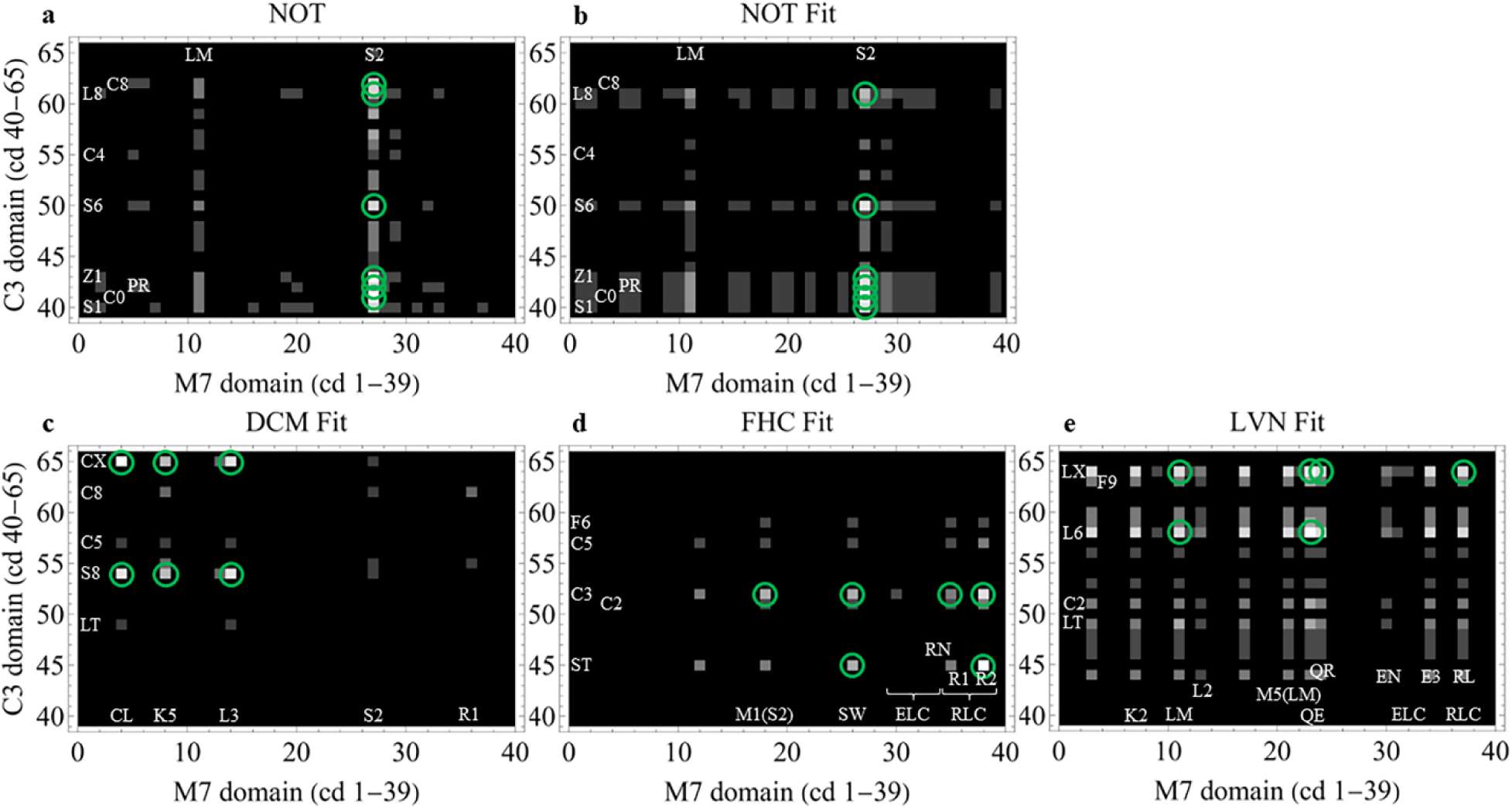
(**panel a**) 2D correlation genetics (2D-CG) map for the NOT phenotype. NOT phenotypes are formed from the 7ddp data set after excluding entries with DCM, FHC, and LVN phenotypes. (**Panels b-e**) 2D-CG fitted maps for NOT, DCM, FHC, and LVN phenotypes.

### 2D-CG map interpretation using thick filament structure basis

The Model 13 neural-symbolic construct introduces thick filament structure parameter *fs* to its neural network module for the βmys/MYBPC3 complex. This symbolic module merges the neural network module with prior knowledge via the human agent (**SI Figure S4**) interpreting: (i) *a priori* 2D-CG maps, (ii) structural inputs from newer cryo-EM [2], cryoelectron tomography [3, 4], and X-ray diffraction data [16], and (iii) other structural information from the literature (see section **Neural-symbolic contraction model**). Diverse inputs promise deeper insight into the role of thick filament structure in disease that is pursued with a thick filament structural basis interpretation of the output 2D-CG maps to deduce coupling strengths for filament structure categories (*sr, dr, xr,* and *gr*, symbolizing, sequestered/off-relaxed, disordered/on-relaxed, other-relaxed structure, and generic-relaxed structure) with the four phenotypes under consideration (DCM, FHC, LVN, NOT). It measures thick filament structure probabilities per disease phenotype.

**Figure 6 panels b-e** indicate fitted 2D-CG maps, constructed as described in METHODS (**Thick filament structure basis**) for NOT, DCM, FHC, and LVN phenotypes best approximating Model 13 2D-CGs (**Figure 6 panel a** and **Figure 2 panels d-f**). Each map (except for DCM) registers modest quantitative differences in co-domain intensities between Model 13 and fitted constructs where the 6 most significant co-domains (encircled) redistribute. All most significant co-domains (in Model 13 and fitted maps) correspond to actual SNV modifications sites (see **Figures 4 and 5**).

The SNV data set described in METHODS consists of 26,055 7ddp entries. It is the basis for quantities summarized in **Tables 1** and **2** below. Data was collected worldwide from people with SNV modified βmys or MYBPC3. Results summarized below reflect properties of this population subset in contrast with a randomly sampled population subset. A randomly sampled population subset might be approximated within the SNV dataset by limiting investigation to phenotypes that have a benign pathogenicity. This option is under investigation.

**Table 1** summarizes filament structure probability for the Model 13 2D-CG maps representing each phenotype. All filament structures contribute substantially to the NOT 2D-CG map (row 4) reflecting its structural heterogeneity. DCM, FHC, and LVN phenotype maps (rows 1-3) have sequestered/off-relaxed (*sr*), disordered/on-relaxed (*dr*), and other-relaxed (*xr*) filament structures predominating, respectively. This contrasts with cardiomyocyte native human myofibrils where *sr* and *dr* state occupancy is ∼42% and 58% [29]. Model 6 2D-CG maps (**Figure 2a-c**) figure in the thick filament structure assignments for the 7ddp dataset (along with other data from the literature, see Cases 1-4 in Results section **Neural-symbolic contraction model**). Their overlap with Model 13 maps (see **Figure 2**) favors this very specific correlation between phenotype and filament structure. **Table 1** re-affirms that the 2D-CG maps reflect a specific correspondence with filament structure induced by SNV’s causing the DCM, FHC, or LVN phenotypes. Comparing the native human myosin model with human SNV data suggests the human disease filament structural specificity reflects an anticipated bias (towards *sr* for DCM or *dr* for FHC). There is no native control for the SNV based studies of thick filament structure. Using the NOT phenotype as a pseudo-control (i.e., assuming without evidence that NOT phenotypes negligibly perturb *sr/dr* equilibrium from native) the DCM SNVs convert most of *dr* and *gr* structures in the control to *sr* while leaving control *xr* unaffected. FHC SNVs convert all spare capacity to *dr*.

**Table 1.** Filament structure probabilities for Model 13 2D-CG maps in. **Figures 2d-2f and 6a estimated using the thick filament structure basis and constrained linear least squares fitting.** Each best model scenario in the solution set of 26 has a unique filament structure basis. Results are averages over model scenarios and errors are standard deviation of the mean for n=26.

| **Table 1: Filament structure probabilities per phenotype specific 2D-CG map** |  |  |  |  |
| --- | --- | --- | --- | --- |
| **2D-CG map** | **SR** | **DR** | **XR** | **GR** |
| 1. DCM (Fig 2d) | 0.85±0.07 | 0.08±0.01 | 0.07±0.07 | 0.00±0.01 |
| 2. FHC (Fig 2e) | 0.00±0.02 | 0.98±0.04 | 0.02±0.02 | 0.00±0.01 |
| 3. LVN (Fig 2f) | 0.03±0.02 | 0.08±0.02 | 0.89±0.02 | 0.00±0.01 |
| 4. NOT (Fig 6a) | 0.51±0.07 | 0.23±0.05 | 0.08±0.04 | 0.18±0.06 |

**Table 2** estimates thick filament structure probabilities for individual DCM, FHC, LVN, and NOT phenotypes (rows 1-4), and, for all phenotypes combined (row 5 labeled ALL). **Table 2** rows 1-4 indicate DCM or FHC disease causing SNVs (but not LVN) contain large fractions of *sr* or *dr* structures with the residual in other structural alternatives represented by the *gr* filament structure. Phenotypes LVN and NOT do not involve other thick filament structures. **Table 2** rows 1-2 indicate DCM or FHC phenotype corresponds to ∼40% or ∼68% *sr* or *dr* filament structure content, respectively, with the remainders in the generic (*gr*) category. Native human ventriculum has 42% or 58% of the relaxed myosins in the *sr* or *dr* structural states [29]. It is unknown if other thick filament structures contribute or even exist for the native myosin.

**Table 2.** Filament structure probabilities for DCM, FHC, LVN, and NOT phenotypes using the thick filament structure basis. Rows 1-4: Contributions to individual phenotypes from the 4 2D-CG maps combined. Each phenotype has a characteristic filament structure probability distribution. Row 5: Filament structure probabilities for all phenotypes combined.

| **Table 2:** Filament structure probabilities per phenotype |  |  |  |  |
| --- | --- | --- | --- | --- |
|  | **SR** | **DR** | **XR** | **GR** |
| 1. DCM | 0.40±0.03 | 0.03±0.03 | 0.02±0.07 | 0.55±0.04 |
| 2. FHC | 0.00±0.01 | 0.68±0.01 | 0.00±0.01 | 0.32±0.01 |
| 3. LVN | 0.01±0.04 | 0.00±0.01 | 0.99±0.05 | 0.00±0.03 |
| 4. NOT | 0.00±0.01 | 0.00±0.01 | 0.01±0.01 | 0.99±0.01 |
| 5. ALL | 0.19±0.04 | 0.48±0.03 | 0.30±0.02 | 0.03±0.01 |

Discrepancies between the Model 13 and fitted 2D-CG maps identify interesting inconsistencies that invite comment. **Figure 6 panel c** shows the DCM Fit 2D-CG map. There minor contributions from MYBPC3 *c8* (co-domaining with βmys *k5*, *s2*, and *r1*) are not reflected in **Figure 2a** indicating the latter’s qualitative mismatch with expectations from the cryo-EM data (see Case 1 in RESULTS). MYBPC3 domain *c8* does not contribute significantly to the Model 13 DCM 2D-CG (**Figure 2d**) despite its explicit introduction into the Case 1 *sr* filament structure category based on cryo-EM data [2]. This discrepancy could be because mavacamten was in the cryo-EM sample preparation. Mavacamten influences coincident-pair domain probability in native human myofibrils [29]. Nonetheless, MYBPC3 *c8* involved co-domains are apparently present in the NOT phenotype 2D-CG map (**Figure 6 panel a**) and plausibly imported to the DCM 2D-CG Fit map by least squares fitting overlap.

### Summary

Models 6 and 13 agree qualitatively on co-domain assignments for the phenotypes (DCM, FHC, and LVN) implying that the general network contraction model (Model 6) and 2D-CG had appropriately assessed the structural basis for these diseases. Now expressed explicitly, using insights from the new structural data [2], co-domains linking βmys sites over multiple myosin dimer crown levels with MYBPC3 C-terminal segment *c4*-*cx* (like Case 1), or, co-domains linking multiple myosin dimer crown levels with MYBPC3 N-terminal segment *st*-*c5* (like Case 2), reflect expectations for relaxed sequestered/off or relaxed disordered/on filament states. Explicit formulation of the filament structure (*fs*) parameter and its SNV influencers with Model 13 identifies intervention targets for deliberately diverting pathogenicity towards benign.

## DISCUSSION

Muscle proteins genetics encode instructions for building a native muscle system and for its impairment by inheritable diseases. An effort to decode the genetics focuses on the βmys/MYBPC3 complex involving the motor (βmys) and one of its regulators (MYBPC3). The complex plays a role in inheritable heart disease phenotypes DCM, FHC, and LVN because it is a locus for their disease causing SNVs.

Model 6 is a general network contraction model decoding βmys/MYBPC3 complex genetics. Its logic emulates the native system to the extent that when the native system is modified by a SNV, the model anticipates the nature (phenotype) and severity (pathogenicity) of functional modification. Model 6 is a feed-forward neural network using known independent inputs: protein residue sequence position, residue substitution, human population, and allele frequency, to surmise two (sometimes unknown) dependent output parameters: phenotype and pathogenicity. The neural network models the causal links between the input SNV characteristics and output phenotype and pathogenicity. It predicts phenotype and pathogenicity assignment given the SNVs characteristics and in this sense decodes βmys/MYBPC3 complex genetics to capture human ventricular muscle functionality.

Genetic decoding identifies crucial phenotype specific inter-protein contacts (co-domains) in the βmys/MYBPC3 complex by using two-dimensional correlation genetics (2D-CG). 2D-CG leverages, a human population genetic divergence phenomena called serial founder effect [12], and the genetic database world-wide surveillance area to locate co-domains in βmys/MYBPC3 [10]. It indicates specific and characteristic co-domain distributions for DCM, FHC, and LVN phenotypes. Phenotype specific co-domain distributions for DCM and FHC diseases suggested they reflect unbalanced thick filament structures, designated here with sequestered/off or disordered/on conformations, that in a native muscle maintain optimum contraction efficiency under changing workloads. Sequestered/off vs disordered/on myosin structures represent energy-conserving vs energy-consuming myosin adaptations [2, 16, 30]. DCM or FHC diseases cause pathological inefficiencies by favoring sequestered/off or disordered/on relaxed myosin conformations contrary to optimal balance [22].

Model 6 decodes the genetic instructions for the contraction mechanism to surmise how SNVs impair its function. Explicit incorporation of thick filament structure, drawing on the sequestered/off vs disordered/on structural concepts, into a new model (Model 13) creates a neural-symbolic hybrid representation for the structural basis of these diseases. The merger accommodates independent filament state data from the literature to be purposefully introduced to the dataset as fulfilled 7ddps constraining neural model optimization. It combines βmys/MYBPC3 interprotein contact sites for all thick filament structures from the Model 6 2D-CG maps [1], thick filament structure data from cryo-EM [2–4], and other data from the literature. The process is illustrated in **SI Figure S4**.

Specific βmys-RLC/MYBPC3 contacts from cryo-EM data raises a question about why 2D-CG maps for DCM causing SNVs (**Figure 2a**) do not indicate co-domains linking RLC with MYBPC3 (**Figure 3a**). Minimal SNVs in RLC for DCM were recorded (at *rn* see **Figure 4**) presumable because RLC SNVs normally cause phenotypes other than DCM. It is noteworthy in this context that βmys/MYBPC3 co-domain contact can appear spontaneously, based on the model prediction of the unassigned phenotype, pathogenicity, or both, for a physical SNV in the unfulfilled part of the database (see **Figure 5**). It is a model generated hypothesis to be tested by future addition of new human genetics data (including its distribution over ethnic human populations) and possibly the application of cryo-EM, or other protein structure determining methods, to the disease impaired assembled muscle components.

Other 2D-CG map analytics use the thick filament structural basis to estimate the structural composition of DCM, FHC, LVN, and NOT phenotypes. It quantitates expected effects of SNV modification on thick filament structure potentially anticipating an excess of disordered/on myosin heads (like in FHC), an excess of sequestered/off myosin heads (like in DCM), or the presence of new structural alternatives not yet identified. Discrepancies between the Model 13 and the fitted 2D-CG maps (compare **Figures 2d and 6c**), the latter created using the thick filament structural basis, identify inconsistencies needing further investigation. For instance, domain de-coupling for CrH/MYBPC3 interactions at *k5/c8*, *r1/(c4 or s8)* and *r1/c8* for the DCM phenotype affected *in vivo* system (**Figure 2d** with data from National Center for Biotechnology Information or NCBI) contrasts with the *in vitro* system in the presence of mavacamten [2] and with the fitted 2D-CG map (**Figure 6c**). The *in vivo* system database has SNVs positioned in domains *k5* and *c8* (**Figure 4**) that should couple if they form a co-domain like that indicated by the *in vitro* system cryo-EM data and the fitted 2D-CG map.

Other proteins in the sarcomere besides βmys/MYBPC3 are implicated in inheritable heart diseases. Actin/βmys, actin/MYBPC3, and actin/thin-filament-regulatory-protein are bi-molecular structures potentially using co-domains to function and where SNVs could cause disfunction. The size and accuracy of the genetic database is the foundation for these new analytical methods. The database inevitably expands over time and more recent genetic data acquisition and deposition into the NCBI database (National Center for Biotechnology Information) have diversified into new human populations (but unwisely eliminated gender from the data set). Likewise, protein structure determining methods applied to the intact muscle system will add physical constraints to the neural-symbolic hybrid contraction model. These trends strengthen model building accuracy within the virtuous cycle *Interpret-revise-repeat* suggested in **SI Figure S4**.

### Conclusion

All machine-learning models start from human-insight. The process described here introduces a workable means to combine new information content from diverse data modalities into an established and evolving model. In this case, it is the timely up-conversion of 6ddps to 7ddps by addition and specification of a new dependent parameter for thick filament structure. Up-conversion is the current and ongoing human-insight aspect in the hybrid model development accomplished with genetic and protein structural data. The goal? A digital twin heart that interprets SNV modifications structurally, discovers disease molecular mechanisms, probes them for likely disease remedies, checks remedy efficacy, and does it without animal models [17].

## METHODS

### SNV data

SNV data retrieval from the National Center for Biotechnology Information (NCBI) is identical to that described previously [1]. SNV dataset from Build 155 released June 16, 2021 is consistent over the present and earlier work on Model 6. SNV reference number (rs#) identifies the affected gene position and one or more alleles corresponding to synonymous and missense variants. Missense variants provide 4 inputs for the SNV defining the protein domain affected (cd), residue substitution (re), population group (po), and allele frequency (af). Associated clinical data gives 2 output parameters, phenotype (*ph*) and pathogenicity (*pa*). Inputs are always known. Outputs are frequently unknown or contradictory among data submitters. Contradictory outputs are decided by consensus when feasible or are designated unknown (*uk*). Input and output parameters form a 6-dimensional data point (6ddp) that is one row in a matrix collecting like data from each SNV in the record. The subset of 6ddps containing unknown output parameters (*ph, pa*, or both) form the unfulfilled 6ddps. The subset of 6ddps containing all known parameters form the fulfilled 6ddps. Application of Model 6 fills in all unknowns forming the unfulfilled/fulfilled 6ddps. All 6ddps are fulfilled in the combined fulfilled and unfulfilled/fulfilled 6ddps matrix. The fulfilled 7ddp dataset is constructed directly from the fulfilled 6ddp dataset as described in **Results: Neural-symbolic contraction model**. All other 7ddps (unfulfilled and unfulfilled/fulfilled) are defined like the analogous 6ddps except Model 13 replaces Model 6.

**SI Figure S2** shows homology (for βmys) and linearized structural models (for βmys and MYBPC3) indicating some mutual binding sites and the locations of most domains. **SI Figure S3** has the complete list of protein domains (*cd*), 2 letter abbreviations, and sequence location. The βmys/MYBPC3 protein complex is made from the four genes, MYH7, MYL2, MYL3, and MYBPC3 having domains from 65 sites and as described [10]. Every SNV in the database has an assigned domain.

Residue substitution (re) refers to the nonsynonymous SNV reference and substituted residue (ref/sub) pairs. Residue abbreviations are the standard 1 letter code. Ref/sub combinations have 420 possibilities for 21 amino acids. Nonsynonymous SNV substitutions for the βmys/MYBPC3 complex in the NCBI database has 147 unique ref/sub pairs.

Human population group and allele frequency fill out the independent parameters in the network. **SI Figure S5** indicates population groups and their 3 letter abbreviations. Allele frequency (af) is a continuous variable in the database on the interval 0 ≤ *af* ≤ 1 for 1 meaning all alleles are substituted by the SNV. These data are divided into two discrete categories of ≤1% (category 0) or >1% (category 1) for this application.

The NCBI SNP database has 17 phenotype data classifications for cardiovascular disease pertaining to βmys and MYBPC3 variants. **SI Figure S1** lists phenotype names and two letter codes. Some phenotypes associate with both βmys and MYBPC3 SNVs. Pathogenicity data classifications include pathogenic (pt), likely pathogenic (lp), benign (be), likely benign (lb), and unknown (uk).

### Models

The Model 6 DAG in **Figure 1** shows the general neural network contraction model described earlier [10]. Inputs {*cd, re, po, af*} affect both outputs {*ph, pa*} while *ph* is causal for *pa*. Model 13 DAG has similar dependencies but involves the new dependent filament structure parameter *fs*. Inputs {*cd, re, po, af*} affect all outputs {*fs, ph, pa*} while *ph* is causal for *pa* and *fs* is causal for both *ph* and *pa*. Input and output parameters for Model 13 form a 7-dimensional data point (7ddp) that is one row in a matrix collecting like data from each SNV in the record. Known values for *fs* are surmised from the 2D-CG representation of results from the application of Model 6 as described in **Results: Neural-symbolic contraction model** and from the literature. The subset of 7ddps containing any combination of 1, 2, or 3 unknown outputs {*fs*, *ph*, *pa*} form the unfulfilled 7ddp matrix. The subset of 7ddps containing all known parameters form the fulfilled 7ddp matrix. The fulfilled 7ddp matrix has rows identical in number and content to the fulfilled 6ddp matrix except for the inclusion of the column containing known filament structure (*fs*).

### Neural network training and validation

**Figure 1** neural networks model structure/function influences from disease and imitate the DAG pathways linking inputs {*cd,re,po,af*} to outputs {*ph,pa*} for Model 6 [9] and outputs {*fs,ph,pa*} for Model 13. Prototype neural networks for both models are shown in the figure. They are tasked with predicting unknown: phenotype and pathogenicity (Model 6) or filament structure, phenotype and pathogenicity (Model 13) from input parameters defining a SNV.

Highest-level optimization implies adapting 1-3 hidden layer depths, and the (1-3) x (input node count) hidden layer widths for both model architectures in **Figure 1**. Each optimization run is called a trial. Trials generate 1-3 best implicit ventriculum contraction model network solutions called general network contraction model scenarios. The process is repeated to accumulate ≥ 20 distinct best scenarios.

The lower-level optimization involves the neural networks populating both model architectures in **Figure 1**. These specific models are trained and validated using the validating dataset composed of all fulfilled 6ddps for Model 6 or 7ddps for Model 13. The training dataset is half of the validating data with members chosen randomly but subject to the constraint that each output is represented in the training dataset except when their representation in the validating dataset is < 2 occurrences. Learnable parameters in the networks are randomly initialized. Weight initializations are normally distributed with zero mean and standard deviation of (1/n)^½^ for n inputs. Bias is initialized to zero. Loss, the number of incorrect predictions from inputs for outputs, is minimized by adjusting learnable parameters in the neural networks using stochastic gradient descent. The validation dataset tests each trained network using a preconfigured loss layer Mathematica routine (CrossEntropyLossLayer) appropriate for output from the softmax layer (**Figure 1**). A preconfigured dropout layer Mathematica routine (**Figure 1**) mitigates overfitting. Overfitting is likewise avoided by terminating training when the validating dataset loss begins to increase following the initial descent during iterative loss minimization. The latter overfitting control is adjusted as appropriate for the model complexity based on form (Models 6 or 13), hidden layer depth and width. The ability to correctly classify SNVs measures the overall suitability of each model scenario. The best 20-30 scenarios form a solution set.

### βmys/MYBPC3 transduction mechanism modeling with Bayes networks

**Figure 1** shows DAGs for the Neural/Bayes network models. Arrows imply a direction for influence hence the domain, residue substitution, population, and allele frequency assignment collectively imply probability for: phenotype and pathogenicity (Model 6) or filament structure, phenotype, and pathogenicity (Model 13). Supplementary Information contains the “All” 6ddp or 7ddp data sets (6ddpdatasetAll.xls or 7ddpdatasetAll.xls each having 26,055 variants) and the “Full” 6ddps or 7ddps (6ddpdatasetFull.xls or 7ddpdatasetFull.xls with 4,092 variants) for complexed βmys/MYBPC3. “All” or “Full” datasets include fulfilled and unfulfilled row entries or fulfilled row entries only. Fulfilled plus unfulfilled/fulfilled datasets (not in **SI**) have unfulfilled entries fulfilled in turn by each of the (20-30) best model scenarios. They define conditional probabilities for the systems in the form of conditional probability tables (CPTs). The product of conditional probabilities on the right defines the joint probability density on the left in,

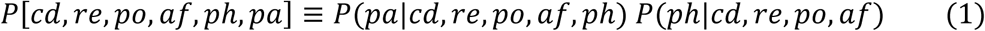

and

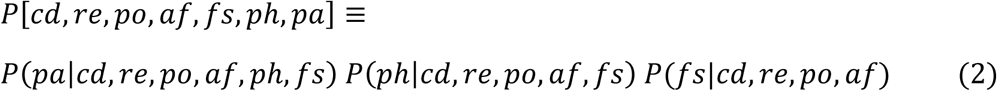

for Models 6 and 13 (eqs. 1 and 2). Calculating SNV probability for domain *i*, population *j*, filament structure *k* (Model 13 only), and phenotype *m* uses joint probability densities,

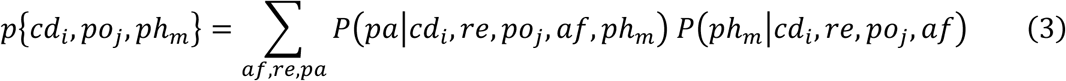

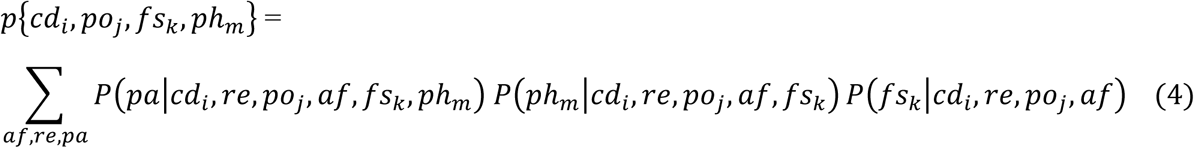

for Models 6 and 13 (eqs. 3 and 4) where summation is over all values for allele frequency, residue substitution, and pathogenicity. Summation over phenotypes defining the broader phenotype categories FHC (*hx, h1, h4*, *h8*), DCM (*ds, d1, dc*), or LVN (*lc, lv, lw*) in eqs. 3 and 4, specifies SNV probability representing each ailment. For example, DCM has probabilities,

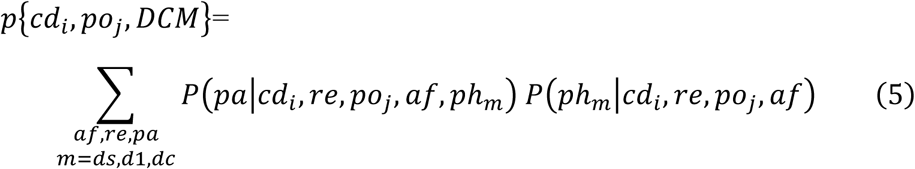

and

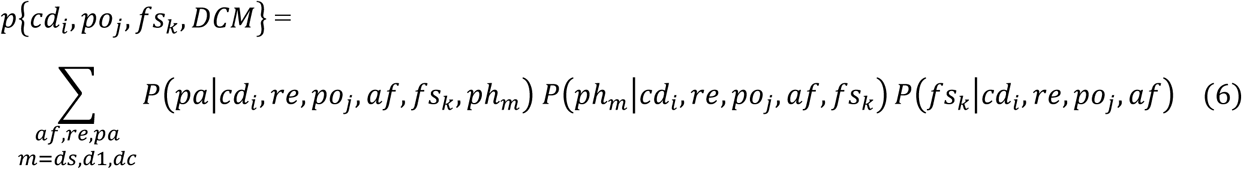

for Models 6 and 13 (eqs. 5 and 6). Model 13 2D-CG computations described below use eq. 6 after summation over filament states as shown in eq. 7,

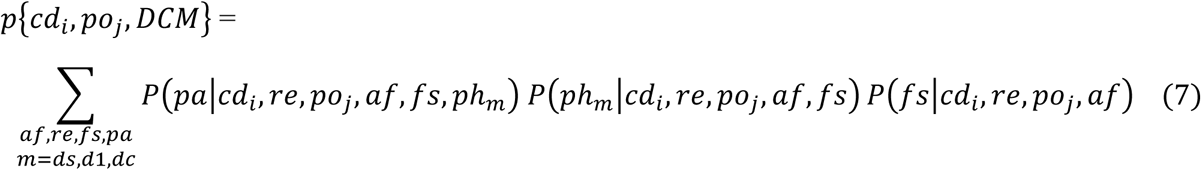

### 2D correlation intensity and significance

SNVs from human population *poj* reside in a protein domain *cdi* with probability given by eq. 5, or 7 for Model 6 or 13, and for each of the broader phenotype categories DCM (shown), FHC, or LVN (these three broader phenotype categories are subsequently indicated collectively with Ω). Eqs. 5 or 7 probabilities populate the synchronous and asynchronous generalized 2D correlation intensities [31] and as described [10],

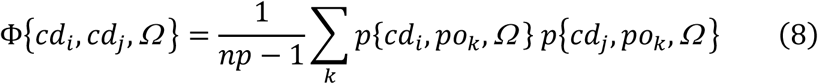

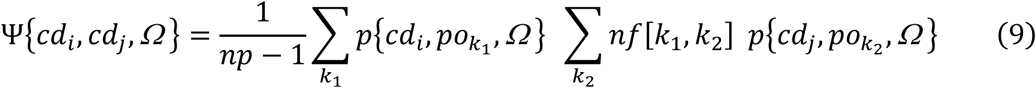

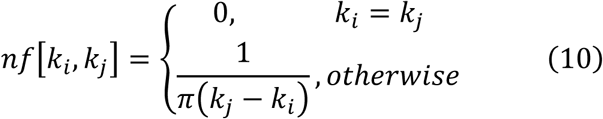

Φ in eq. 8 gives synchronous, and Ψ in eq. 9 asynchronous, 2D correlation intensities for *np* the number of populations represented by *po* and collective phenotype categories Ω = DCM, FHC, or LVN. The 2D synchronous intensity map will identify domain pairs whose SNV probabilities correlate or anti-correlate for identical populations while the 2D asynchronous intensity map identifies the leading and lagging co-domain member over populations ordered by their genetic divergence. The most significant combined synchronous and asynchronous co-domain interaction cross-correlate probability amplitudes are the largest elements in the array,

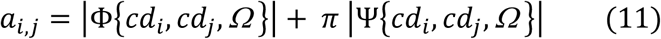

for Φ and Ψ from eqs. 8 and 9, *cdi* and *cdj* the SNV containing functional domains in βmys and MYBPC3, and Ω the phenotype categories DCM, FHC, or LVN. Array element *ai,j* is the co-domain cross-correlate given by the product of probabilities for SNVs in each domain member of the co-domain weighted by synchronous or asynchronous coupling to human populations. Quantity π multiplying the asynchronous cross-correlates (Ψ) in eq. 11 balances weighting for synchronous (Φ) with nearest neighbor population asynchronous cross-correlates. Elements *ai,*j are ≥ 0. They are combined with −*ai,j* and their distribution visualized in a histogram. A normal distribution approximates the result. Absolute distance from the mean expressed in multiples of standard deviation indicates a measure of co-domain cross-correlate amplitude significance. Co-domain cross-correlate amplitude significance increases with distance from amplitude mean.

Data obtained from protein domain SNV probability correlations represented in 2D plots (usually called 2D-CG maps) show amplitude (eq. 11) for βmys (abbreviated M7) on the abscissa and MYBPC3 (C3) domains on the ordinate, in the order given in SI **Figure S3**. Domains are discrete entities hence the 2D-CG maps resemble a pixelated image with 8-bit grayscale representing intensity. Each pixel in a map corresponds to a co-domain pair with inter-protein co-domain correlation strength indicated by intensity. Maps thus constructed are obtained for each of the (20-30) best model scenarios in the solution set.

### Thick filament structure basis

The set of 2D-CG maps dependent solely on filament structures (*sr*, *dr*, *xr*, *gr*) and phenotypes (DCM, FHC, LVN, NOT) define a 16-component basis that covers the disease space. Equation 6 gives the probability construct needed to populate synchronous and asynchronous generalized 2D correlation intensities (the disease space), like those introduced with eqs. 8-10, but retaining explicit filament state (*fs*) dependence as below,

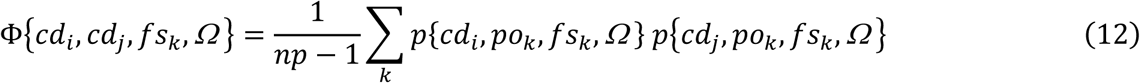

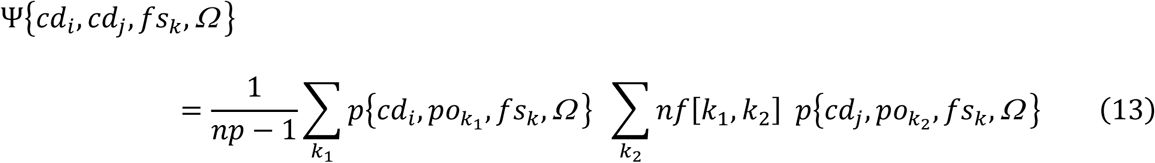

And with nf[ki, kj] from eq. 10. Quantities express synchronous (eq. 12) and asynchronous (eq. 13) map intensities that define basis maps spanning the filament-structure/phenotype space occupied by the Model 13 2D-CG maps (**Figures 2d-2f** and **Figure 6a**). Basis maps thus constructed are computed from CPT’s representing each of the (20-30) best model scenarios in the solution set.

Constrained linear least-squares fitting gives amplitudes, constrained to be ≥ 0, for best fitting the Model 13 2D-CG maps. Best fitted maps compare favorably with observations, compare **Figures 2d-2f and 6a** with **Figures 6c-6e and 6b**, implying the relative contribution of basis filament structures in each phenotype. It provides the methodology to pursue the stated aim to deduce quantitative coupling strengths for the filament structure categories for the phenotypes under consideration.

### Population genetic divergence proxy

Worldwide human genetic divergence is attributed to migration from a single origin in East Africa based on the serial founder effect [12, 32, 33]. Serial founder effect explains observed linear divergence decrease with human migration distance over the earth’s surface. Migration distance is therefore a good proxy for genetic divergence variation. Estimates for migration distances on earth’s surface are calculated exactly as described [10]. Distance estimates (and genetic divergence variation) from the fitted line for populations in **SI Figure S6** are equally spaced over the Population Index parameter as needed for the co-domain 2D-CG formalism [31].

## SUPPLEMENTARY INFORMATION

Supplementary information consists of a document file with **Figures S1-S6**, and four raw data sets for complex βmys/MYPBC3. Data sets are in files 6ddpdatasetAll.xls, 6ddpdatasetFull.xls, 7ddpdatasetAll.xls, and 7ddpdatasetFull.xls for Model 6 (6ddps) and Model 13 (7ddps) cases. Data set names containing “All” or “Full” include fulfilled and unfulfilled entries or fulfilled entries only. Fulfilled plus unfulfilled/fulfilled datasets vary with each of the 20-30 scenarios in the solution dataset and are not included. They are available from the author.

## Availability of data and materials

All data generated or analyzed in the study are included in the published article and its supplementary information files. All computer coding is in Mathematica (Wolfram Research, Champaign, IL) and available by request from the author.

## Acknowledgement

Thanks to Katalin Ajtai for scientific discussion and critical review of the manuscript.

## Author contribution

TPB is the sole author and responsible for the content of the paper. The author read and approved the final manuscript.

## Funding

This research did not receive any specific grant from funding agencies in the public, commercial, or not-for-profit sectors.

## Notes

https://www.ncbi.nlm.nih.gov/

### Competing Interest Statement

The authors have declared no competing interest.

