## Supplementary Information for "Neural-symbolic hybrid model for myosin complex in cardiac ventriculum decodes structural bases for inheritable heart disease from its genetic encoding"

Thomas P. Burghardt

Department of Biochemistry and Molecular Biology

200 First St. SW

Mayo Clinic Rochester

Rochester, MN 55905

https://orcid.org/0000-0003-3119-4074

19 April 2024

Revised 31 August 2024

| code | phenotypes | index |
|---|---|---|
| d1 | Dilated cardiomyopathy 1A (MyBPC3) | 1 |
| dc | Familial dilated cardiomyopathy | 2 |
| dm | Myopathy, distal, 1 (MYH7) | 3 |
| ds | Dilated cardiomyopathy 1S (MYH7) | 4 |
| h1 | Familial hypertrophic cardiomyopathy 1 (MYH7) | 5 |
| h4 | Familial hypertrophic cardiomyopathy 4 (MYBPC3) | 6 |
| h8 | Familial hypertrophic cardiomyopathy 8 (MYL3) | 7 |
| hb | Hyaline body myopathy (MYH7), Myosin storage myopathy, Myopathy, myosin storage, autosomal recessive | 8 |
| hc | Familial hypertrophic cardiomyopathy, Primary familial hypertrophic cardiomyopathy, Hypertrophic cardiomyopathy, Increased left ventricular wall thickness, Left ventricular hypertrophy, Concentric hypertrophic cardiomyopathy | 9 |
| hd | Cardiomyopathy, hypertrophic, midventricular, digenic (MYLK2) | 10 |
| hx | Familial hypertrophic cardiomyopathy 10 (MYL2) | 11 |
| lc | Left ventricular noncompaction 10 (MYBPC3) | 12 |
| lv | Left ventricular noncompaction cardiomyopathy, Left ventricular noncompaction | 13 |
| lw | Left ventricular noncompaction 5 (MYH7) | 14 |
| m7 | MYH7-Related Disorders | 15 |
| rc | Familial isolated restrictive cardiomyopathy (MYBPC3, MYL3) | 16 |
| uk | conflicting-interpretations-of-phenotype, not specified, uncertain-significance, NA | 17 |

**Figure S1.** Phenotype (*ph*), 2 letter codes, and descriptive names from the database. Each phenotype associates with SNVs falling within the genes comprising human βmys/MYBPC3 (MYH7, MYL2, MYL3, and MYBPC3). Conflicting-interpretation-of-phenotype (code *uk*) implies no consensus phenotype from the database. The DCM, FHC, and LVN phenotype categories refer to combined sub-phenotypes {*ds, d1, dc*}, {*hx, h1, h4, h8*}, and {*lc, lv, lw*}, respectively.

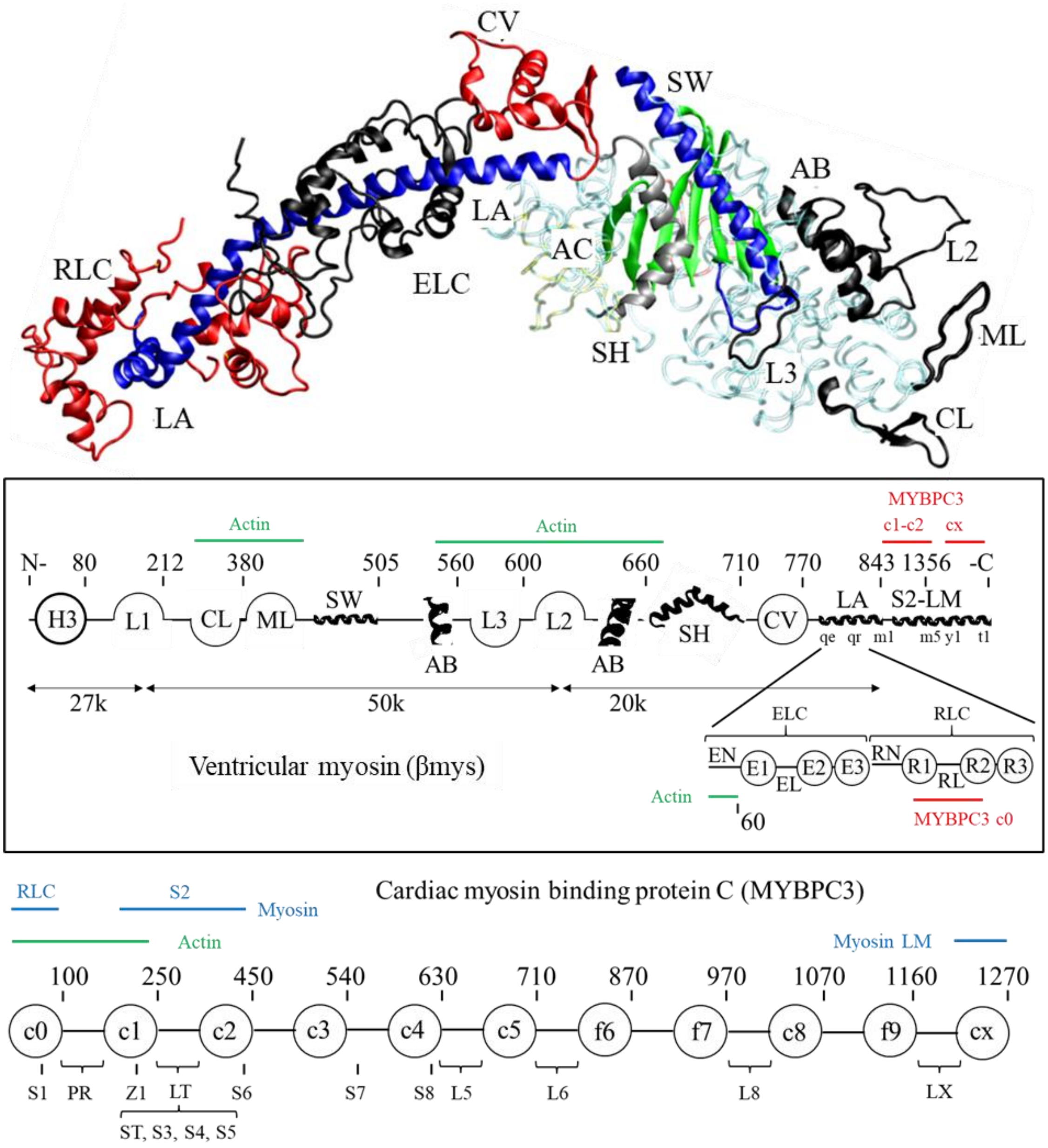


**Figure S2.** Homology and linearized models for the βmys structure (top and middle) and the linearized model representing MYBPC3 (bottom), identifying many of the domains listed in **SI Figure S3**. The linearized myosin diagram (middle diagram inside box) does not indicate active site (*ac*), OM binding site (*om*), mesa (*me*), and blocked head/converter binding site (*bh*) because they occupy multiple regions in the linearized representation. Myosin light chains appear below the heavy chain. Several known MYBPC3 and actin binding sites are indicated in red and green above the heavy chain and below the light chains. The MYBPC3 diagram (bottom) has 8 Ig-like domains (denoted with prefix *c*) and 3 fibronectin-like domains (prefix *f*). Serine phosphorylation sites: S1 (Ser47), ST (Ser275), S3 (Ser284), S4 (Ser304), S5 (Ser311), S6 (Ser427), S7 (Ser550), and a threonine phosphorylation site S8 (Thr607) are indicated below the chain. Domain linkers of interest include the proline rich linker (PR) and linker 2 (LT) containing a regulatory site. LT, L5, L6, and L8 are probable swivels. Z1 inside *c1* is a zinc binding site. Myosin RLC, S2, and LM (in blue) and actin binding (in green) sites on MYBPC3 are indicated above the linearized model.

| domain (cd) | seq | index |
|---|---|---|
| M7 actin binding (ab) | <525-563, 647-655> | 1 |
| M7 active site (ac) | <114-125, 167-178, 244-253, 260-267, 453-465, 666-673> | 2 |
| M7 Blocked Head/Converter Binding (bh) | various 295-397 | 3 |
| M7 C-loop (cl) | 359-377 | 4 |
| M7 Converter (cv) | 711-768 | 5 |
| M7 SH3 (h3) | 28-78 | 6 |
| M7 20k (k2) | 634-843 | 7 |
| M7 50k (k5) | 209-633 | 8 |
| M7 27k (k7) | Nterm-208 | 9 |
| M7 Lever-arm (la) | 769-843 | 10 |
| M7 LMM (lm) | 1357-Cterm | 11 |
| M7 Loop 1 (l1) | 202-212 | 12 |
| M7 Loop 2 (l2) | 622-646 | 13 |
| M7 Loop 3 (l3) | 564-576 | 14 |
| M7 MESA (me) | various 168-664 | 15 |
| M7 Myopathy-loop (ml) | 400-414 | 16 |
| M7 MESA-trail (mr) | various 172-917 | 17 |
| M7 S2-C1 mybpc3 (m1) | 844-857 | 18 |
| M7 S2-LT mybpc3 (m2) | 924-936 | 19 |
| M7 S2-C2 mybpc3 (m3) | 858-870 | 20 |
| M7 LMM-Cx (m5) | 1554-1581 | 21 |
| M7 OM binding (om) | various 91-712 | 22 |
| M7 IQ ELC (qe) | 788-798 | 23 |
| M7 IQ RLC (qr) | 814-824 | 24 |
| M7 SH1/SH2 hinge (sh) | 683-710 | 25 |
| M7 Switch 2 helix (sw) | 466-505 | 26 |
| M7 Subfragment 2 (s2) | 844-1356 | 27 |
| M7 LMM-titin (t1) | 1815-1831 | 28 |
| M7 LMM-myomesin (y1) | 1506-1674 | 29 |
| M7 N-term ELC (en) | Nterm-60 | 30 |
| M7 EF1 ELC (e1) | 49-86 | 31 |
| M7 linker ELC (el) | 87-127 | 32 |
| M7 EF2 ELC (e2) | 128-163 | 33 |
| M7 EF3 ELC (e3) | 164-Cterm | 34 |
| M7 N-term RLC (rn) | Nterm-23 | 35 |
| M7 EF1 RLC (r1) | 24-59 | 36 |
| M7 linker RLC (rl) | 60-94 | 37 |
| M7 EF2 RLC (r2) | 95-129 | 38 |
| M7 EF3 RLC (r3) | 130-166 | 39 |
| C3 phospho Ser 1 (s1) | 43-51 | 40 |
| C3 c0-Ig like (c0) | 1-101 | 41 |
| C3 proline rich (pr) | 102-152 | 42 |
| C3 zinc site 1 (z1) | <208, 210, 223, 225> | 43 |
| C3 c1-Ig like (c1) | 153-256 | 44 |
| C3 phospho Ser 2 (st) | 271-279 | 45 |
| C3 phospho Ser 3 (s3) | 280-288 | 46 |
| C3 phospho Ser 4 (s4) | 300-307 | 47 |
| C3 phospho Ser 5 (s5) | 308-315 | 48 |
| C3 linker c1-c2 (lt) | 257-361 | 49 |
| C3 phospho Ser 6 (s6) | 423-431 | 50 |
| C3 c2-Ig like (c2) | 362-452 | 51 |
| C3 c3-Ig like (c3) | 453-543 | 52 |
| C3 phospho Ser 7 (s7) | 546-554 | 53 |
| C3 phospho Thr 8 (s8) | 603-611 | 54 |
| C3 c4-Ig like (c4) | 544-633 | 55 |
| C3 linker c4-c5 (l5) | 634-644 | 56 |
| C3 c5-Ig like (c5) | 645-711 | 57 |
| C3 linker c5-f6 (l6) | 712-743 | 58 |
| C3 c6-Fibronectin (f6) | 744-870 | 59 |
| C3 c7-Fibronectin (f7) | 871-967 | 60 |
| C3 linker c7-c8 (l8) | 968-970 | 61 |
| C3 c8-Ig like (c8) | 971-1065 | 62 |
| C3 c9-Fibronectin (f9) | 1066-1163 | 63 |
| C3 linker c9-c10 (lx) | 1164-1180 | 64 |
| C3 c10-Ig like (cx) | 1181-1274 | 65 |

**Figure S3.** Protein domain names and two letter codes (*cd*), protein sequence assignment (*seq*), and indexed numbering for domains in the βmys/MYBPC3 complex. Domain names begin with M7 or C3 designating origin from βmys or MYBPC3, respectively. Myosin default domains 27k, 50k, and 20k refer to the molecular weights for tryptic proteolytic fragments from cleavage of the myosin heavy chain (MYH7) sequence in Loop 1 at the active site, and Loop 2 in the actin binding site [1]. OM binding (*om*, index number 22) is the binding site for Omecamtiv Mecarbil [2]. Several domains are identified in **SI Figure S2** structures.

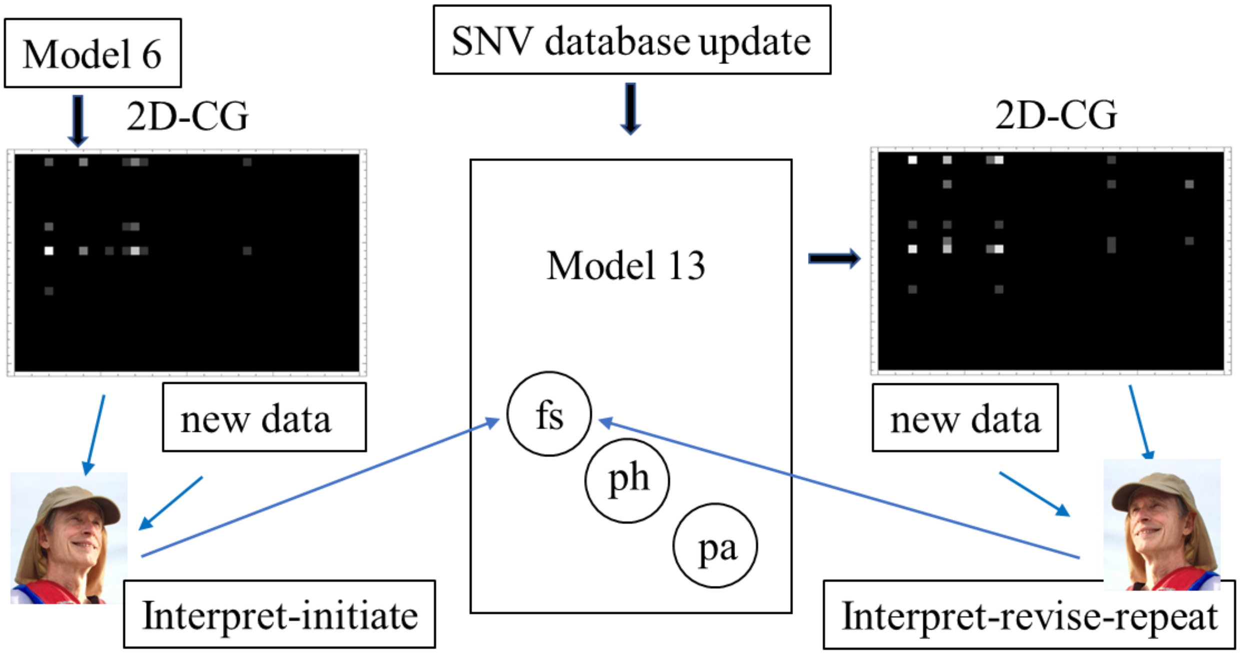


**Figure S4**. Neural-symbolic contraction model origin and adaptation. Model 6 is bootstrapped into Model 13 by a human agent (author's photograph) interpreting the Model 6 2D-CG map and data from the literature for initial assignment of filament structure (*fs*) parameters. Model 13 neural network optimization and creation of the new 2D-CG map follows. Additional interpret-revise cycles occur whenever new structural data is available or with revision of Model 13. Model revision could require introduction of a new parameter and involve bootstrapping for their initial assignments.

| code | population | index |
|---|---|---|
| ACP | ACPOP | 1 |
| AFM | African American | 2 |
| AFR | African | 3 |
| AMR | American | 4 |
| ASI | Asian | 5 |
| ASJ | Ashkenazi Jewish | 6 |
| CEA | CentralAmerican | 7 |
| CRM | Chinese | 8 |
| CUB | Cuban | 9 |
| DAN | Danish | 10 |
| DOM | Dominican | 11 |
| EAS | East Asian | 12 |
| EST | Estonian | 13 |
| EUA | European American | 14 |
| EUR | European | 15 |
| FIN | Finnish from FINRISK project | 16 |
| GON | Genome of the Netherlands | 17 |
| JPN | JAPANESE | 18 |
| KOR | Korean | 19 |
| LA1 | Latin American 1 | 20 |
| LA2 | Latin American 2 | 21 |
| MDE | Middle_Est | 22 |
| MEX | Mexican | 23 |
| NAM | NativeAmerican | 24 |
| NHI | NativeHawaiian | 25 |
| OCE | Oceania | 26 |
| OTH | Other | 27 |
| PCC | PARENT AND CHILD COHORT | 28 |
| PRI | PuertoRican | 29 |
| SAS | SouthAsian | 30 |
| SOA | SouthAmerican | 31 |
| SPC | Spanish controls | 32 |
| TWC | TWIN COHORT | 33 |

**Figure S5**. Human population codes and descriptions from 1000Genomes, gnomAD-Genomes, TopMed, and other studies in the NCBI database including: ACPOP (ACP) whole-genome sequenced control population study from Västerbotten County in northern Sweden; Dominican (DOM) Dominican Republic; Latin American 1 (LA1) Latin American individuals with Afro-Caribbean ancestry; Latin American 2 (LA2) Latin American individuals with mostly European and Native American Ancestry; Other (OTH) missense SNVs where a population category is not indicated; Parent and Child Cohort (PCC) UK10K Avon Longitudinal Study of Parents and Children Variants; Spanish controls (SPC) Medical Genome Project healthy controls from Spanish population; Twin Cohort (TWC) UK10K Department of Twin Research and Genetic Epidemiology twin registry of 11,000 identical and non-identical twins between the ages of 16 and 85 years.

| Population Sequence | Code | Distance from Addis Ababa km | Line km | Limits km | Outlier | Index |
|---|---|---|---|---|---|---|
| Middle_Est | MDE | 3934 | 3934 | {3518, 5936} | | 1 |
| African | AFR | 4526 | 4526 | {1190, 7002} | | 2 |
| Ashkenazi Jewish | ASJ | 5118 | 5118 | {2480, 8149} | | 3 |
| European | EUR | 5710 | 5710 | {5424, 8149} | | 4 |
| Genome of the Netherlands | GON | 6302 | 6302 | {6000, 9039} | | 5 |
| Danish | DAN | 6894 | 6894 | {6122, 9182} | | 6 |
| PARENT AND CHILD COHORT | PCC | 7486 | 7486 | {6215, 9406} | | 7 |
| European American | EUA | 8078 | 8078 | {6355, 9622} | | 8 |
| SouthAsian | SAS | 8670 | 8670 | {5929, 9815} | | 9 |
| Spanish controls | SPC | 9262 | 9262 | {6703, 10013} | | 10 |
| Asian | ASI | 9854 | 9854 | {6558, 10754} | | 11 |
| American | AMR | 10446 | 10446 | {6687, 10829} | | 12 |
| Oceania | OCE | 11038 | 11038 | {7411, 11531} | | 13 |
| African American | AFM | 11630 | 11630 | {5610, 12193} | | 14 |
| Latin American 1 | LA1 | 12222 | 12222 | {5282, 12662} | | 15 |
| Cuban | CUB | 12768 | 12814 | {7661, 12768} | * | 16 |
| Estonian | EST | 13406 | 13406 | {7862, 14229} | | 17 |
| PuertoRican | PRI | 13998 | 13998 | {8609, 14509} | | 18 |
| Finnish from FINRISK project | FIN | 14590 | 14590 | {7869, 14640} | | 19 |
| East Asian | EAS | 15118 | 15182 | {9572, 15118} | * | 20 |
| Korean | KOR | 15774 | 15774 | {9548, 16416} | | 21 |
| Dominican | DOM | 16365 | 16365 | {10288, 17577} | | 22 |
| SouthAmerican | SOA | 16957 | 16957 | {11171, 18362} | | 23 |
| Latin American 2 | LA2 | 17549 | 17549 | {12425, 19697} | | 24 |
| CentralAmerican | CEA | 18141 | 18141 | {12755, 21166} | | 25 |
| Mexican | MEX | 18733 | 18733 | {13784, 22886} | | 26 |
| NativeAmerican | NAM | 19325 | 19325 | {15912, 28074} | | 27 |
| NativeHawaiian | NHI | 19940 | 19917 | {19940, 31181} | * | 28 |
| Column 1 | 2 | 3 | 4 | 5 | 6 | 7 |

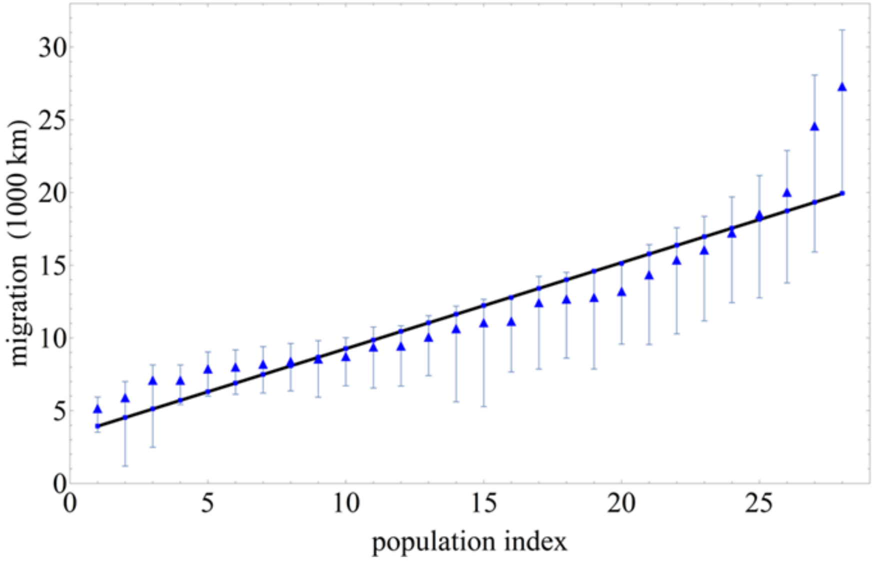


**Figure S6**. Migration distance vs human population index (from **SI Figure S5**) and pertaining to complex βmys/MYBPC3 in tabular (top) and graphical (bottom) forms. **Top**. Populations (column 1) listed are in a linear relationship with migration distance (column 3). The migration distance is a proxy for genetic divergence variation where divergence decreases with distance. Fitted line (column 4), computed as described [3], closely follows migration distances (column 3) and falls within limits (column 5) except for outliers identified with the asterisk in column 6. The population index (column 7) identifies populations in graphical presentations like that in the bottom panel. **Bottom**. Thick black line indicates a best estimate for the linear relationship of listed populations with migration distance. Light blue vertical bars at each triangle show minimum limits needed to fulfill the linear estimate as described in the text. Blue triangles indicate migration distances falling within the blue vertical bars best fitted by the blue line.
